# Type I fatty acid synthase (FAS) trapped in the octanoyl-bound state

**DOI:** 10.1101/747683

**Authors:** Alexander Rittner, Karthik S. Paithankar, Aaron Himmler, Martin Grininger

## Abstract

*De novo* fatty acid biosynthesis in humans is accomplished by a multidomain protein, the type I fatty acid synthase (FAS). Although ubiquitously expressed in all tissues, fatty acid synthesis is not essential in normal healthy cells due to sufficient supply with fatty acids by the diet. However, FAS is overexpressed in cancer cells and correlates with tumor malignancy, which makes FAS an attractive selective therapeutic target in tumorigenesis. Herein, we present a crystal structure of the condensing part of murine FAS, highly homologous to human FAS, with octanoyl moieties covalently bound to the transferase (MAT) and the condensation (KS) domain. The MAT domain binds the octanoyl moiety in a novel (unique) conformation, which reflects the pronounced conformational dynamics of the substrate binding site responsible for the MAT substrate promiscuity. In contrast, the KS binding pocket just subtly adapts to the octanoyl moiety upon substrate binding. Besides the rigid domain structure, we found a positive cooperative effect in the substrate binding of the KS domain by a comprehensive enzyme kinetic study. These structural and mechanistic findings contribute significantly to our understanding of the mode of action of FAS and may guide future rational inhibitor designs.

**Highlights:**

- The X-ray structure of the KS-MAT didomain of murine type I FAS is presented in an octanoyl-bound state.
- Multiple conformations of the MAT domain and a dynamic active site pocket explain substrate promiscuity.
- The rigid domain structure and minor structural changes upon acylation are in line with the strict substrate specificity of the KS domain.
- Enzyme kinetics reveals cooperativity in the KS-mediated transacylation step.

## Introduction

Fatty acids are essential molecules in most living cells, serving as key compounds of cell membranes, as energy supply in the metabolism, as secondary messengers in signaling pathways or as covalent modifications to recruit proteins to membranes. They can either be obtained directly from the diet or are synthesized *de novo* by fatty acid synthases (FASs) from simple building blocks in repeating cyclic reactions. Although the chemistry of fatty acid synthesis is largely conserved across all kingdoms of life, the structural organization of the participating enzymes differs dramatically. FAS complexes occurring in plants, bacteria and in mitochondria, known as the type II, perform biosynthesis by a series of monofunctional separate enzymes (Beld et al., 2015; Chen et al., 2018; White et al., 2005). In contrast, the CMN group bacteria (*Corynebacterium*, *Mycobacterium*, and *Nocardia*), fungi and higher eukaryotes utilize type I FASs that integrate all enzymatic functions into large macromolecular assemblies (Grininger, 2014; Heil et al., 2019; Herbst et al., 2018; Maier et al., 2010). Fungal and CMN-bacterial FASs form up to 2.7-MDa α_6_β_6_-heterododecameric barrel-like structures (Boehringer et al., 2013; Elad et al., 2018; Johansson et al., 2008; Leibundgut et al., 2007; Lomakin et al., 2007). The animal FASs, including human FAS, emerge from a separate evolutionary development and consist of two polypeptide chains assembling into a 540-kDa intertwined “X-shaped” homodimer (Maier et al., 2008).

In animals, including human beings, fatty acid biosynthesis commences with the transfer of an acetyl moiety from acetyl-coenzyme A (CoA) to the terminal thiol of the phosphopantetheine arm of the acyl carrier protein (ACP) domain catalyzed by the malonyl-/acetyltransferase (MAT) domain (Figure 1A) (Smith and Tsai, 2007). After being passed on to the active site cysteine of the β-ketoacyl synthase (KS) domain, a malonyl moiety is loaded on the free ACP domain in a second MAT-mediated transfer reaction. Upon delivery of the malonyl moiety, the KS domain catalyzes a decarboxylative Claisen condensation reaction in which the KS-bound acetyl and the ACP-bound malonyl moieties combine to an ACP-bound β-ketoacyl intermediate. Subsequently, the β-keto group is sequentially modified by three processing domains, the ketoreductase (KR), the dehydratase (DH) and the enoylreductase (ER) using NADPH as a reducing agent. Typically, fatty acid synthesis runs seven cycles to deliver a fully reduced ACP-bound acyl chain of 16 carbon atoms, which is eventually released as palmitic acid by hydrolysis via the thioesterase (TE) domain (Figure 1A).

**Figure 1:**
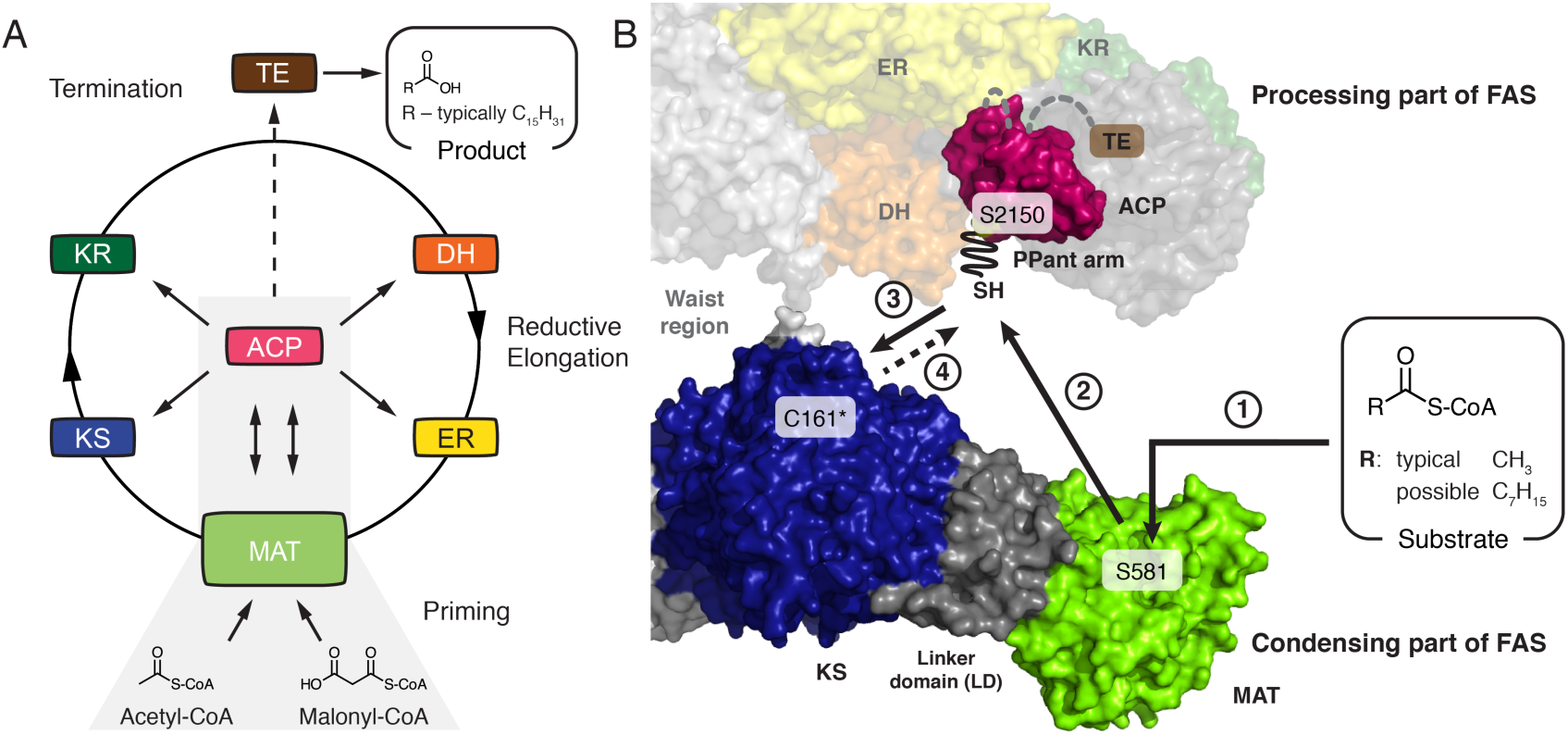
Priming of animal fatty acid synthesis. In a first step, the substrate is selected by the MAT domain and transferred to the ACP domain (2. step) from where it is passed on to the KS domain (3. step). Important active site residues are highlighted and C161 is marked with an asterisk. Crystal structure of porcine FAS (PDB code 2vz9) and NMR structure of rat ACP (PDB code 2png) are depicted in surface representation (Maier et al., 2008; Ploskoń et al., 2008). Domains of one protomer of FAS homodimer are colored. Domain nomenclature: MAT, malonyl-/acetyltransferase; ACP, acyl carrier protein; KS, ketosynthase; KR, ketoreductase; DH, dehydratase; ER, enoylreductase; TE, thioesterase; PPant arm expand fully in the figure.

The key domains in fatty acid synthesis are MAT, KS and ACP, responsible for the selection of substrates, C-C bond formation and substrate shuttling, respectively (**Figure 1B**). The MAT domain of murine type I FAS shows broad substrate specificity and facilitates the synthesis of methyl-branched, odd numbered and functionalized fatty acids by alternative substrate selection (Buckner et al., 1978; Rittner et al., 2019; Rittner et al., 2018; Smith and Stern, 1983). On the contrary, the KS domain possesses a strict specificity for saturated acyl moieties with a low acceptance of β-keto groups to guarantee biosynthesis of saturated fatty acids *in vivo* (Witkowski et al., 1997). The ACP domain is loaded by the MAT and ACP does not impose substrate specificity in this step. However, the interaction of ACP with other domains is necessary for the progress of synthesis and the specificity of this interaction can affect the product formation (Dodge et al., 2019; Rossini et al., 2018; Sztain et al., 2019).

Although FASs are ubiquitously expressed in all tissues, *de novo* biosynthesis of fatty acids occurs at low levels as the demand is usually met by the diet (Semenkovich et al., 1995; Uhlén et al., 2015). Despite adequate nutritional lipid supply, the FAS gene is overexpressed under pathological conditions, being associated with diseases like diabetes, obesity and cancer, and upregulation of FAS correlates with tumor malignancy (Gansler et al., 1997; Khandekar et al., 2011; Kuhajda, 2006; Menendez et al., 2009; Nguyen et al., 2010; Rashid et al., 1997). FAS has emerged as a very promising therapeutic target in tumorigenesis, because pharmacological FAS inhibitors induce tumor cell death by apoptosis whereas normal cells are resistant (Pandey et al., 2012). To date, several inhibitors, like cerulenin, GSK2194069 and orlistat, have been identified or developed that target the KS, KR and TE domain, respectively (Hardwicke et al., 2014; Pandey et al., 2012; Pemble et al., 2007). Remarkably, the compound TVB-2640 recently entered phase 2 clinical trials showing promising results in combinatorial anti-cancer therapies (Buckley et al., 2017; Dean et al., 2016).

Herein, we report the crystal structure of the murine KS-MAT didomain at 2.7 Å resolution with octanoyl moieties covalently bound in the KS and the MAT active sites. By comparison with structures of domains in apo-form as well as with the malonyl-bound MAT domain, we analyze structure-function relationships and correlate the conformational variability of the individual domains with their substrate specificities. Furthermore, by applying a continuous fluorometric assay, we reveal detailed mechanistic insight into the cooperative behavior of the KS-mediated transacylation reaction. The results of this study provide new insights into the key processes of substrate loading and condensation in fatty acid synthesis and foster the development and optimization of inhibitors with potential antineoplastic properties.

## Results

### Crystal structure of the KS-MAT didomain with bound octanoyl moieties

In order to gain structural insights into the molecular basis for the substrate ambiguity of the MAT domain and the strict substrate specificity of the KS domain of murine type I FAS, we aimed at trapping both KS and MAT domains in the octanoyl-bound enzyme state. Following an established protocol (Rittner et al., 2018), the purified murine KS-MAT didomain, sharing 87 % sequence identity to the condensing part of human FAS (Pappenberger et al., 2010), was crystallized and crystals were soaked with octanoyl-CoA (Figure S1). X-ray diffraction data were collected to a resolution of 2.7 Å, and the resulting structural model refined to R/R_free_ of 0.18/0.23 (Table 1). The asymmetric unit contains four polypeptide chains (A-D) arranged as two biological dimers interacting via the cleft between the KS and the linker domain (LD) (**Figure 2A**). In all four chains, the KS domain is modified with an octanoyl moiety yielding the octanoyl-enzyme covalent complex. Furthermore, in one polypeptide chain (chain D), an octanoyl group is covalently bound to S581 of the MAT domain and an additional octanoyl-CoA is non-covalently trapped in the MAT binding tunnel (Figure 2B). This finding confirms previous data showing that octanoyl-CoA can prime murine type I fatty acid synthesis and that the MAT domain can catalyze the transfer of octanoyl moieties (Rittner et al., 2019; Rittner et al., 2018).

**Figure 2:**
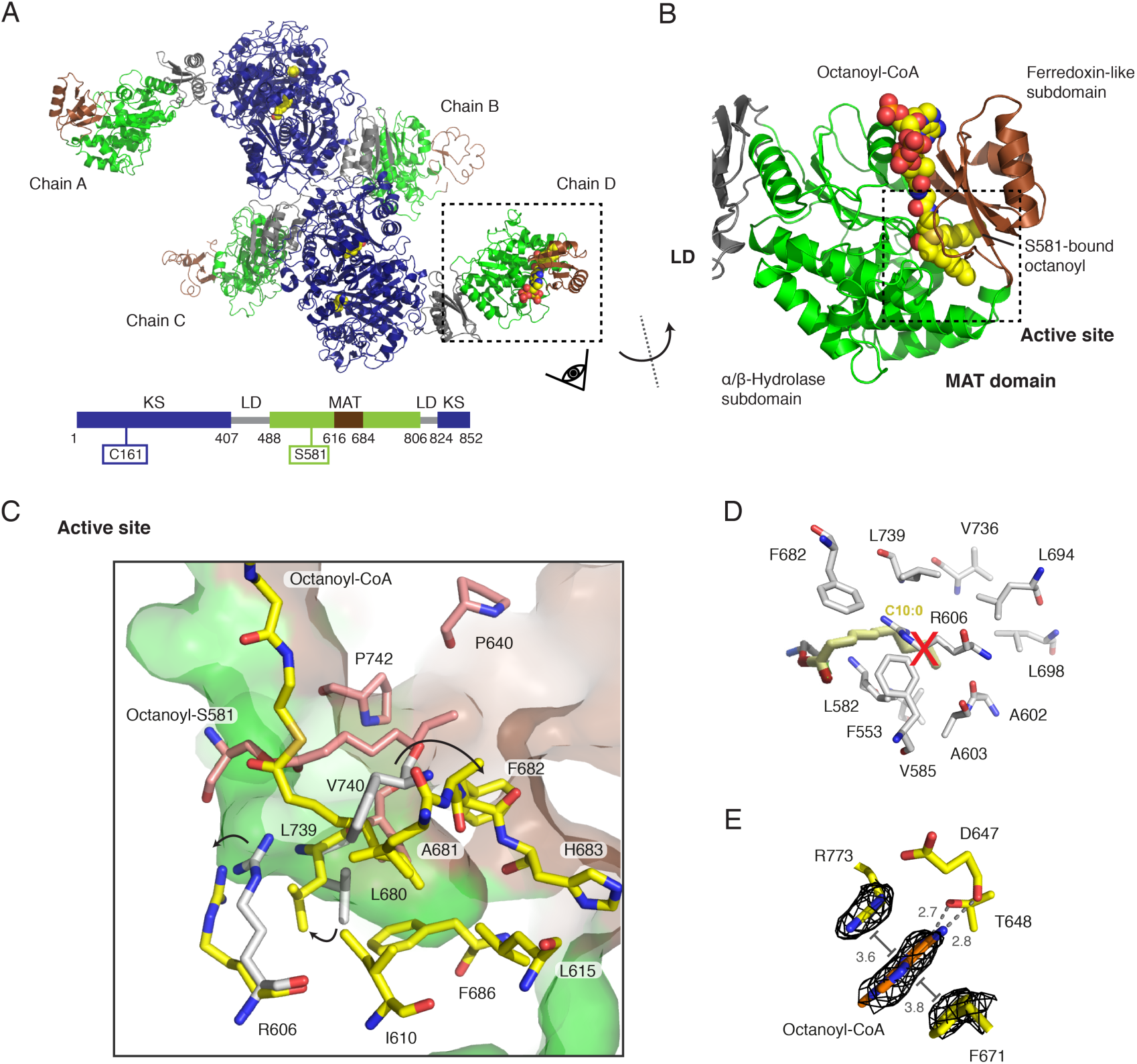
Octanoyl-loaded MAT domain. (A) Polypeptide chains in the unit cell with bound octanoyl moieties in yellow in sphere representation. Domains and folds are coloured as depicted in the attached cartoon. (B) Zoom into the MAT domain in chain D. The active site is embedded in a cleft between the α/β-hydrolase (green) and the ferredoxin-like (brown) subdomains. The active site serine (S581) was found in an octanoyl-bound state with an additional octanoyl-CoA molecule non-covalently attached to the active site tunnel (see Figure S2 for FEM (Afonine et al., 2015) and Polder maps (Liebschner et al., 2017)). (C) Zoom into the MAT active site. Residues interacting with the serine-bound octanoyl chain and the octanoyl CoA are coloured in red and yellow, respectively. Movements of select residues upon binding of an octanoyl moiety in comparison with the human MAT structure (grey) are indicated by arrows. (D) Orientation of a decanoyl chain as reported by Bunkoczi *et al*..(Bunkoczi et al., 2009). Atomic coordinated originate from decanoyl chain computationally modelled into the human MAT variant R606A (pdb code: 2jfd). Postulated interacting residues of the human MAT domain are shown in grey. (E) Binding site of the nucleobase of the CoA moiety at the MAT surface. The adenine is coordinated via hydrogen bonding by with residues D647 and T648 and via π-stacking and π-cation interactions with residues F671 and R773 were identified.

**Table 1.** Data collection and refinement statistics.

|  |  |
| --- | --- |
|  | Wild-type soaked with octanoyl-CoA |
| Data Collection |  |
| Space group | C222<sub>1</sub> |
| Cell dimensions |  |
| a, b, c (Å) | 147.4 354.0 218.5 |
| α, β, γ (°) | 90 90 90 |
| Resolution (Å) | 50-2.7 (2.75 – 2.7) |
| No. of reflections | 2195612(110844) |
| R<sub>meas</sub> | 0.16(2.4) |
| I/σI | 13.9(1.4) |
| CC<sub>1/2</sub> | 0.99(0.62) |
| Completeness (%) | 98.2(98.8) |
| Redundancy | 14.3(14.5) |
| Refinement |  |
| R<sub>work</sub>/R<sub>free</sub> (%) | 18.4(23.2) |
| No. of unique reflections | 145548 |
| Average B factors (Å<sup>2</sup>) |  |
| Protein | 78.9 |
| Wilson *B* factor | 58.4 |
| RMSD from ideality |  |
| Bond length (Å) | 0.007 |
| Bond angles (°) | 1.5 |
| Ramachandran statistics |  |
| Favored regions (%) | 93.43 |
| Allowed regions (%) | 5.05 |
| Outliers (%) | 1.52 |

Highest resolution shell is shown in parantheses.

### General description of the octanoyl-bound MAT domain

The MAT domain engages fatty acid synthesis in selecting the CoA-ester substrates for product assembly. It is located at the edge of the condensing part of animal FAS and inserted into the KS fold via the linker domain (LD). The exposed position and the utilization of only 8.4 % of the solvent-accessible area for domain-domain interactions reflect a high structural independence from the FAS fold. The substrate-binding pocket is formed by a cleft between the α/β-hydrolase and the ferredoxin-like subdomains and extends to the active site located in the center of the domain. The function of the MAT domain is to shuttle acyl moieties via the active serine between CoA-ester substrates and the ACP domain following a ping-pong bi-bi mechanism. The catalytic key residues S581 and H683 form a catalytic dyad with S581 acting as nucleophile and H683 serving in acid-base catalysis (Paiva et al., 2018). The nucleophilicity of S581 is enhanced by a helix dipole-moment due to its positioning within a strand-turn-helix motif termed the nucleophilic elbow (Hol, 1985). A key residue for the bifunctional role of MAT is R606 located in helix 7, which interacts with the carboxylic group of extender substrates. The absence of the guanidinium group leads to altered substrate specificity (Rangan and Smith, 1997; Rittner et al., 2018).

Upon soaking protein crystals with octanoyl-CoA, we found that S581 in chain D is covalently modified with an octanoyl moiety and, furthermore, a molecule octanoyl-CoA is non-covalently attached to the substrate binding pocket (**Figure 2B**). The placement of both ligands was based on the feature-enhanced map (FEM) and later validated by a Polder map (Figure S2) (Afonine et al., 2015; Liebschner et al., 2017). The octanoyl chain of octanoyl-S581 is located in a tunnel between the two subdomains created between residues P640, F682, V740 and P742 and extends to the protein’s surface (**Figure 2C**). The octanoyl group of octanoyl-CoA points towards helix 10 of the α/β-hydrolase fold in a substrate binding pocket between the subdomains created by residues I610, L615, L680, A681, F682, H683, F686 and L739. Rangan and Smith (1997) reported that the R606A variant from rat FAS showed increased turnover rates for the transfer of octanoyl moieties (Rangan and Smith, 1997). Based on this, Bunkoczi and co-workers (Bunkoczi et al., 2009) placed a decanoyl chain in the human R606A-mutated MAT binding site by a simulated docking experiment and concluded that space for longer acyl chains is created in the mutant due to the absence of the side chain of residue R606. In our structure, the positions of the octanoyl chains are slightly different to that of the computationally docked decanoyl chain. Upon binding the octanoyl chain, the side chains of residues R606 and L680 rotate to form an extended binding cavity that can accommodate the larger substrate. This feature is likely responsible for the high transacylation rate of murine MAT for octanoyl moieties and may not be present in rat MAT according to docking data (Bunkoczi et al., 2009). Again deviating from rat FAS data (Rangan and Smith, 1997), the murine MAT loses the capability for efficient transacylation of the octanoyl moiety upon mutation of R606 to alanine (Table 2).

**Table 2.** Kinetic analysis of the transacylation reaction with octanoyl-CoA at a fixed acceptor concentration of 60 µM ACP (n = 4)

| Substrate | $K_m^{app}$ ($\mu M$) | | | $k_{cat}^{app}$ ($s^{-1}$) | | | $k_{cat}/K_m$ ($M^{-1} s^{-1}$) | | | | | Hydrolysis rate $10^{-3}$ ($s^{-1}$) | | |
| --- | --- | --- | --- | --- | --- | --- | --- | --- | --- | --- | --- | --- | --- | --- |
| Wildtype* | 0.7 | ± | 0.3 | 4.1 | ± | 0.3 | 5.6 | ± | 2.2 | × | $10^6$ | 6.1 | ± | 1.0 |
| R606A variant | 4.9 | ± | 0.7 | 0.037 | ± | 0.008 | 7.6 | ± | 2.0 | × | $10^3$ | 2.3 | ± | 3.4 |

\*Previously published (Rittner et al., 2018).

Intriguingly, in the MAT-octanoyl-CoA complex, the nucleobase of octanoyl-CoA is bound at a specific position between the two subdomains with the adenine stacked between side chains of F671 and R773. Besides the well-known π-stacking between aromatic rings, also π-cation interactions between arginines and aromatic rings are known (Flocco and Mowbray, 1994). Additionally, two hydrogen bonds are formed between amines of the purine ring to both the side chain hydroxyl group of T648 and the backbone carbonyl group of D647.

### Implications on MAT subdomain dynamics from crystal structures

Variations in the relative positioning of the ferredoxin-like subdomain were reported in previous crystal structures, but a correlation of subdomain mobility to the substrate ambiguity of the domain could not yet be drawn. In chain D of MAT-octanoyl-CoA complex, the MAT domain was found in a unique conformational state. Keeping the α/β-hydrolase part of the domain (backbone atoms (BB) of D488–D611 and D685-D806) as a reference, a superposition was performed with the apo-structure in chain A, the malonyl-bound structure (PDB code 5my0; chain D) and the human KS-MAT (PDB code 3hhd; chain A). The α/β hydrolase domain superimposes very well in all the four models with RMSDs (BB) to the MAT-octanoyl-CoA complex of 0.6 (Chain A), 0.8 (Malonyl-bound chain D), and 0.6 (human chain D) Å, respectively (**Figure 3A**). Largest differences are found in the relative positioning of the ferredoxin-like subdomain with local shifts of up to 7.3 Å between corresponding residues. As also the ferredoxin-like subdomains (BB of 618–674) of all four models themselves superimpose well with RMSDs (BB) between 0.4-0.8 Å, these results clearly illustrate that the ferredoxin-like subdomain describes a rigid-body movement.

**Figure 3:**
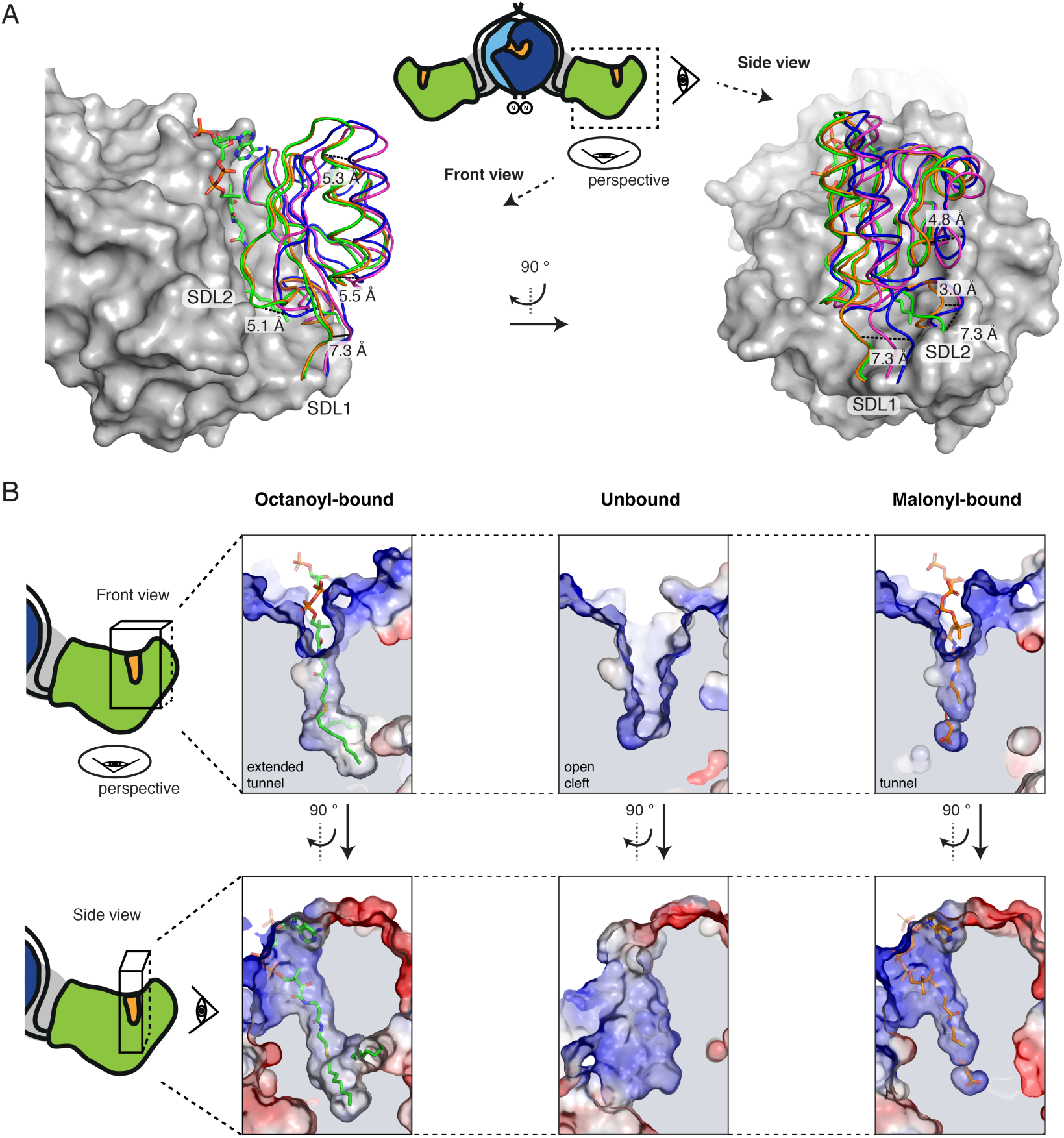
Conformational variability of the MAT active site. (A) α/β-Hydrolase fold centrered superposition (BB of residues 488–613 and 685-806) of four MAT domains in different acyl-bound states. Chain A (blue) and chain D (green) from the octanoyl-CoA soaked crystal (PDB code 6rop), malonyl-bound (orange) (PDB code 5my0; chain D) and apo human MAT (purple) (PDB code 3hhd; chain A) were used. The α/β-hydrolase subdomain is shown in surface depiction and the ferredoxin-like fold in cartoon loops. Selected distances between corresponding residues indicate the mobility of the subdomains with a relative movement of the ferredoxin-like fold of up to 7.3 Å. (B) Different active site and entry tunnel shapes upon substrate binding. Surface depictions of active sites of chain D (left panel) and A (middle) from the octanoyl-bound structure and chain D (right panel) from the malonyl-bound structure are shown in two perspectives. Surfaces are shown in surface electrostatic representation calculated with PyMOL and shown in default coloring with positive potential depicted in blue and negative potential in red. Views as in (A).

When the static X-ray structural information is subjected to a TLS (Translation, Libration and Screw) refinement, the derived anisotropic displacement parameters imply a rotational movement describing the opening and closing of the active site cleft to allow binding of diverse substrates (Figure S5) (Winn et al., 2001). In order to determine residues contributing most to the positional variability of the ferredoxin-like and the α/β-hydrolase subdomains, we plotted main-chain torsion angles φ and ψ (Ramachandran plot) for the MAT domains of the various structural models. The plot identifies residues A613 and H614 as well as H683 and S684 as undergoing significant changes in main-chain torsion angles (Figure S4). Both sites are the hinges of two subdomain linkers, termed SDL1 (612-617) and SDL2 (675-684), allowing movements of SDL1 and SDL2 of about 7.3 Å and 5.1 Å, respectively (Figure S3A). The positional and conformational variability of the subdomain linkers allows changes in the relative orientation of the subdomains and in the geometry of the active site cleft for the accommodation of chemically and structurally diverse CoA esters (Rittner et al., 2018) (**Figure 3B**).

In addition to the overall dynamics of the MAT fold, the residue R606, responsible for holding the carboxyl group of extender substrates, shows high positional variability in the MAT structural models. The high degree of rotational freedom of the side chain originates likely from the specific property of animal MAT in featuring a phenylalanine at a position (F553, murine MAT numbering), which is otherwise occupied by a conserved glutamine. As shown previously, F553 significantly diminishes the coordination of the R606 side chain by hydrogen-bonding (Rittner et al., 2018). In the octanoyl-bound structure, we could identify a third rotameric state of R606, in addition to the ones found in apo- and malonyl-bound state (Figure S6), which demonstrates that the adaptation of the domain to different substrates is closely connected to the rotational variability of this residue.

### Structure of the KS domain in an acylated state

The KS domain forms dimers in type I FAS systems, and contributes the largest area (about 2580 Å^2^; see Table S1 for more information) to the overall dimerization interface of animal FAS. The KS domain belongs to the thiolase-superfamily and exhibits the characteristic topology of alternating layers of α-helices and β-sheets (called α/β/α/β/α sandwich motif) (**Figure 4A**). A small vestibule in lateral orientation to the two-fold axis of the condensing part forms the entry to the active site, which is comprised of the active cysteine (C161) as well as two histidine (H293, H331) residues, termed the catalytic triad. The substrate binding tunnel further extends towards the dimer interface, where it merges with the tunnel of the protomer at the two-fold axis (**Figure 4B**).

**Figure 4:**
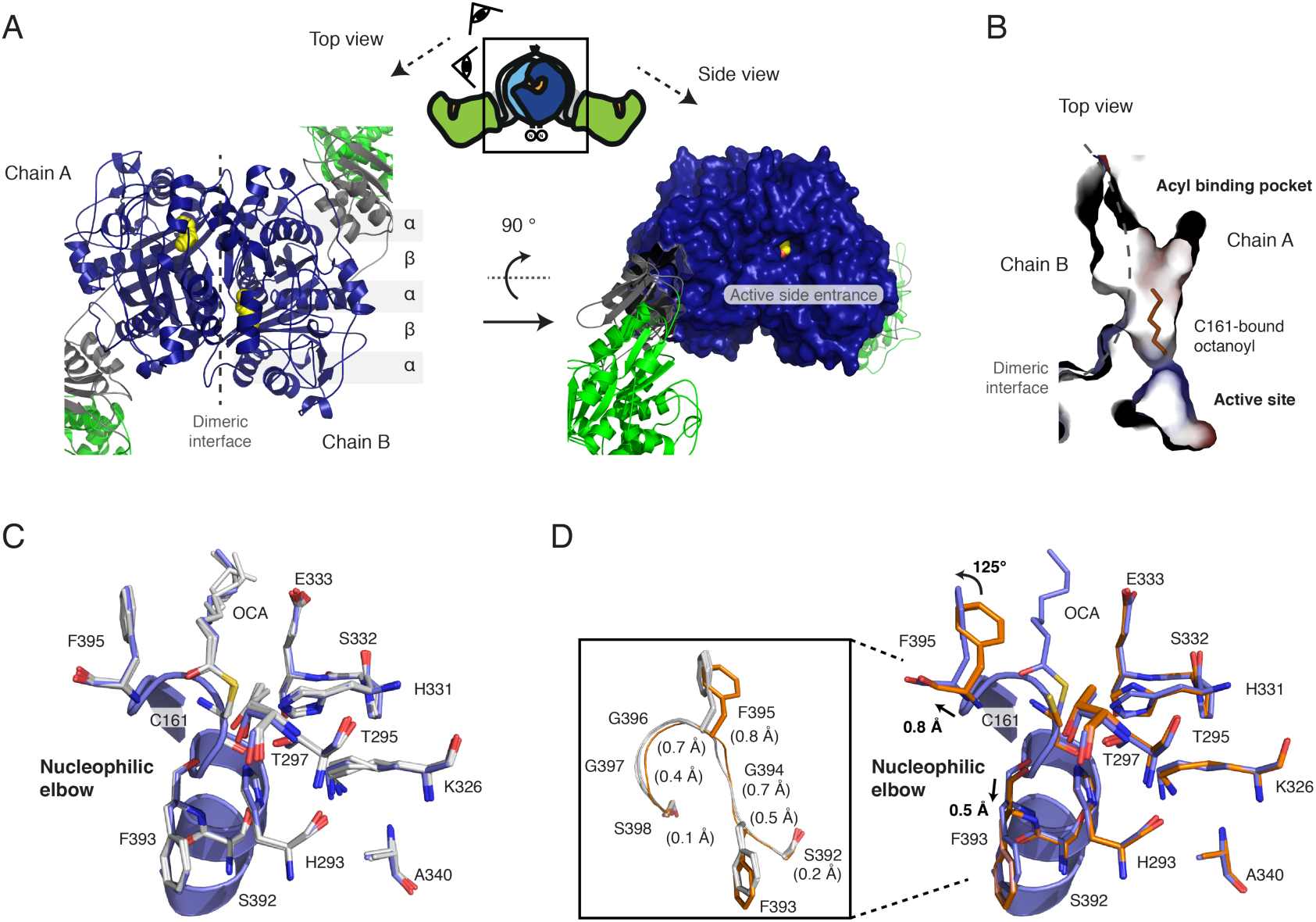
Octanoyl loaded KS domain. (A) Top view on the dimeric KS domain in cartoon depiction showing the topology of the α/β/α/β/α sandwich motif (left panel). A surface depiction of the KS domain in side view highlights the active site entrance. Color codes as in Figure 2A is used with the bound octanoyl chain shown in yellow in sphere representation (right panel). (B) Active site and acyl binding cavity of the KS domain. In addition to the substrate binding cavity between the dimer interface, a small side chamber is visible in the monomer. The binding cavity is shown with surfaces colored in electrostatic potential (colored as in Figure 3). (C) Active site of KS showing important residues for catalysis, reported for homologous KAS I (FabB) (Olsen et al., 2001). Three chains (B-D) with bound octanoyl moieties were aligned to chain A (blue) by a KS based superposition (BB of residues 1–407 and 824–852). All residues adopt essentially the same conformation with some variability in the terminal carbon atoms of the octanoyl chain. (D) A similar KS based superposition was performed with the four apo-KS domains (orange; PDB code 5my0) and the octanoyl-bound chain A (blue). Upon octanoyl binding, the individual residues of the stretch 393-397 are shifted by 0.4-0.8 Å (highlighted in the inlet). Furthermore, the side chain of F395 is rotated by approximately 125°.

Overall all the four polypeptide chains of the asymmetric unit align very well onto one another when a KS domain based superposition is performed (RMSDs (BB) of about 0.20-0.25 Å over the residue ranges 1–407 and 824– 852) (**Figure 4C**). All four active sites in the KS domains are modified with octanoyl moieties at residue C161. The position and conformation of all active site residues are essentially identical. Only the bound octanoyl chain shows positional variability in the terminal carbon atoms due to an unconstrained rotational freedom of the single bonds. Taking also into account the overall low B-factors observed in the KS part of the crystal structure, this data indicates a relatively low degree of flexibility within the KS domain.

As observed for S581 in the MAT domain, the active cysteine C161 is positioned in a nucleophilic elbow. Here, the positive dipole-moment of the α-helix decreases the p*K*_a_ value of the thiol group leading to the increased nucleophilicity of the sulfur. In a recent computational study, it was reported that the thiol group of the active cysteine is readily deprotonated under physiological condition. This implies a role of H331 in acting as a general acid in catalysis, which is different to acid-base catalysis performed by the active histidine of MAT domains (Lee and Engels, 2014). Such a role of H331 is confirmed by our structural data, as nitrogen (ND1) accepts hydrogen bonds from backbone amides of P332 (3.4–3.6 Å) and E333 (3.0–3.4 Å), whereas the protonated nitrogen (NE2) of H331 is in hydrogen bond distance (3.2–3.4 Å) to the sulfur of the thioester bond at residue C161.

Furthermore, the bound octanoyl chains allow localization of the oxyanion hole, which is created by backbone amides of residues C161 and F395. In all chains, the carbonyl’s oxygen of the thioester is in hydrogen bond distance to the corresponding amides of F395 (2.9–3.1 Å) and slightly further apart from backbone amides of C161 (3.1–3.4 Å).

Next, we were interested in conformational changes induced by the loading of an octanoyl chain in comparison to the unbound state. Therefore, the four unbound murine KS domains of the unit cell (PDB code 5my0) were aligned to octanoyl-bound chain A (serving as the representative chain) in a KS domain based superposition (residues 1–407 plus 824–852) (**Figure 4D**). Again, overall RMSDs (BB) were small (0.2-0.3 Å), but the superposition revealed distinct differences in the positions and side chain conformations of some residues. Most prominently, the stretch of residues FGFGG (residues 393-397) is slightly shifted and reorganized upon binding of the octanoyl moiety. This results in the displacement of F395 by 0.8 Å (between corresponding carbonα atoms) plus the rotation of the side chain by approximately 125°. F395 in the rotamer position of the unbound state clashes with the octanoyl chain in the bound state implying that the rearrangement of F395 is necessary to accommodate substrates.

FabB (KAS I) and FabF (KAS II), both elongating β-ketoacyl-ACP synthases of the bacterial type II fatty acid synthesis, have been well-characterized in their three-dimensional structure in an octanoyl-bound state and in a dodecanoyl-bound state, respectively (see e.g. (Olsen et al., 2001; von Wettstein-Knowles et al., 2006; Wang et al., 2006). KS domain based superpositions of FabB and FabF to chain A (BB residues 1–407) show that the active sites display an overall identical architecture implying a conserved catalytic mechanism for the type I KS domain (Figure 5A). An exception is a glutamate residue, which is conserved in CHH class structures (E342 and E349 in FabB and FabF, respectively) and is thought to participate in catalysis by stabilizing a water or cation molecule (Olsen et al., 2001; von Wettstein-Knowles et al., 2006) This acidic residue is exchanged by an alanine in the type I KS domain (A340) excluding an equivalent role in animal FAS.

**Figure 5:**
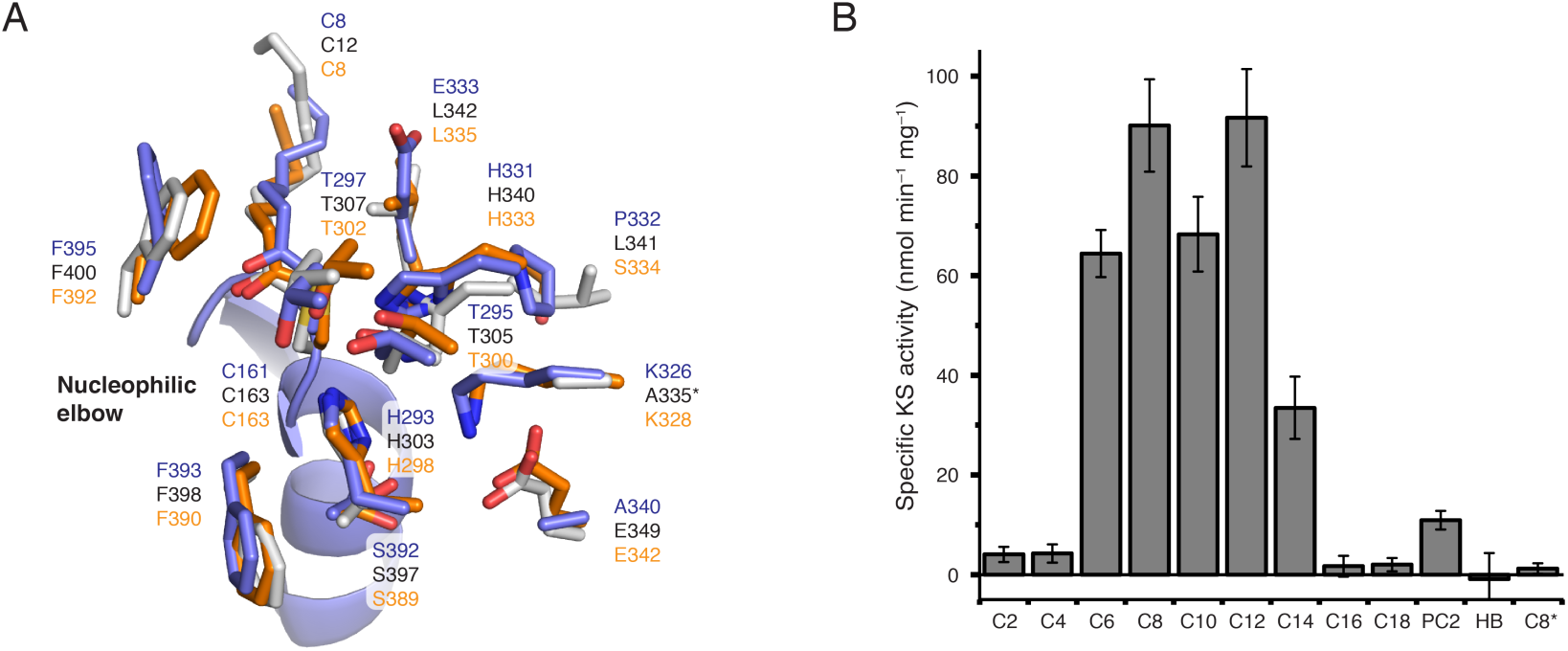
Chain-length specificity of KS and comparison of the KS domain with FabB and FabF from *E.* coli. (A) Comparison of important active site residues of the murine type I KS domain (chain A; blue) with FabB (orange; PDB code 2bui) and FabF (grey; PDB code 2gfy) from *E*. *coli* (von Wettstein-Knowles et al., 2006; Wang et al., 2006). All three proteins were solved in the acyl bound state and *E. coli* proteins were aligned to chain A (blue) by a KS based superposition (BB of residues 1–407). The asterisk indicates that a variant of FabF was crystalized possessing a K335A mutation. (B) Chain-length specific KS-mediated transacylation activity. The specific KS-mediated activity was determined at fixed substrate (500 µM) and holo-ACP (75 µM) concentrations using the αKGDH-assay. The asterisk indicates usage of variant KS^C161G^MAT^S581A^ as negative control. Abbreviations refer to acyl-CoA esters with different chain lengths and PC2 and HB refer to phenylacetyl-CoA and hydroxybutyryl-CoA, respectively.

### Specificity of the KS domain for saturated acyl chains

The first step in the KS-mediated Claisen condensation is the transacylation of an acyl-moiety from acyl-ACP to the active site cysteine (**Figure 1**). Considering the similarity of this step to the MAT-mediated transacylation, we aimed at using the continuous fluorometric assay, originally established for transferase analysis (Molnos et al., 2003; Rittner et al., 2018), to investigate the substrate specificity of the KS domain. In doing so, we have constructed KS-MAT^S581A^ for specific KS read-out and the double knockout mutant KS^C161G^-MAT^S581A^ as a control. All didomain constructs proved to be stable, which was validated by size exclusion chromatography profiles and by melting temperatures obtained in a thermal shift assay (Figure S7). We determined KS-mediated turnover rates from various acyl-CoA esters to a separated, standalone holo-ACP at fixed substrates concentrations (**Figure 5B**). The experiment generally confirmed the results from Witkowski *et al*. of turnover rates increasing with acyl chain-length until maximum rates are reached for octanoyl-CoA and dodecanoyl-CoA (C12-CoA), and decreasing rates with chain-length above C12-CoA (Witkowski et al., 1997). The high value for C12-CoA is not consistent with previous results for the rat homolog and seems to be a specific feature of the heterologously expressed murine KS domain. The specificity of the transfer of acyl-moieties was confirmed with the KS^C161G^-MAT^S581A^ double knockout mutant (Figure 5B).

Further, the substrate specificity of the KS domain was probed with two non-cognate acyl-CoA substrates. While the hydroxybutyryl-CoA, mimicking the intermediate after an initial reduction by the KR-domain, was not accepted as substrate, the non-canonical compound phenylacetyl-CoA was transferred with a reasonable rate. The latter result confirms our previous data that this substrate can also serve as a priming substrate for fatty acid synthesis (Rittner et al., 2019).

### The KS-mediated transacylation shows kinetic cooperativity

To gain insight into the enzymatic properties of the KS domains, the absolute kinetic parameters for the KS-mediated transacylation from acyl-CoA esters to the ACP domain (as a standalone protein) were determined by the assay described before. The KS-mediated transacylation reaction follows a ping-pong bi-bi mechanism with a covalently bound acyl-enzyme intermediate and can be described with the general equation 1 that is based on standard Michaelis-Menten kinetics (Copeland, 2005). In order to determine absolute kinetic parameters, we have used this equation to globally fit two series of response curves with octanoyl-CoA and myristoyl-CoA at five or six different fixed ACP concentrations (see Methods section). The global fit could only moderately describe the dependence of the apparent turnover rates in respect of the individual ACP concentrations and disclosed systematic deviations in the response curves at low and high substrate concentrations (see Figure S8).

For a better description of the sigmoidal shape of the individual plots, and considering the dimeric nature of the KS domain, we included a Hill coefficient for both substrate concentrations (CoA-Ester and ACP) as described in equation 2. This new fit function clearly delineates both data series without imposing any parameter constraints (see **Figure 6**). The absolute kinetic constants (*K’*) and turnover numbers (*k*_cat_) for octanoyl-CoA and myristoyl-CoA are 139 ± 16 µM (*K’*), 0.09 ± 0.007 s^−1^ (*k*_cat_) and 111 ± 8 µM, 0.05 ± 0.002 s^−1^, respectively. These values lead to specificity constants (*k*_cat_/*K’*) in the range of 10^2^–10^3^ s^−1^ M^−1^, which indicate rather inefficient priming of the KS domain by CoA-esters. The *K’*_ACP_ of the standalone ACP was with 16 ± 1.7 µM higher for the transfer of octanoyl moieties than for myristoyl moieties (8 ± 0.8 µM). The calculated Hill coefficients were between 1.7 and 2 for both substrates indicative of a positive cooperativity of the KS domains of a dimeric unit during the transacylation reaction.

**Figure 6:**
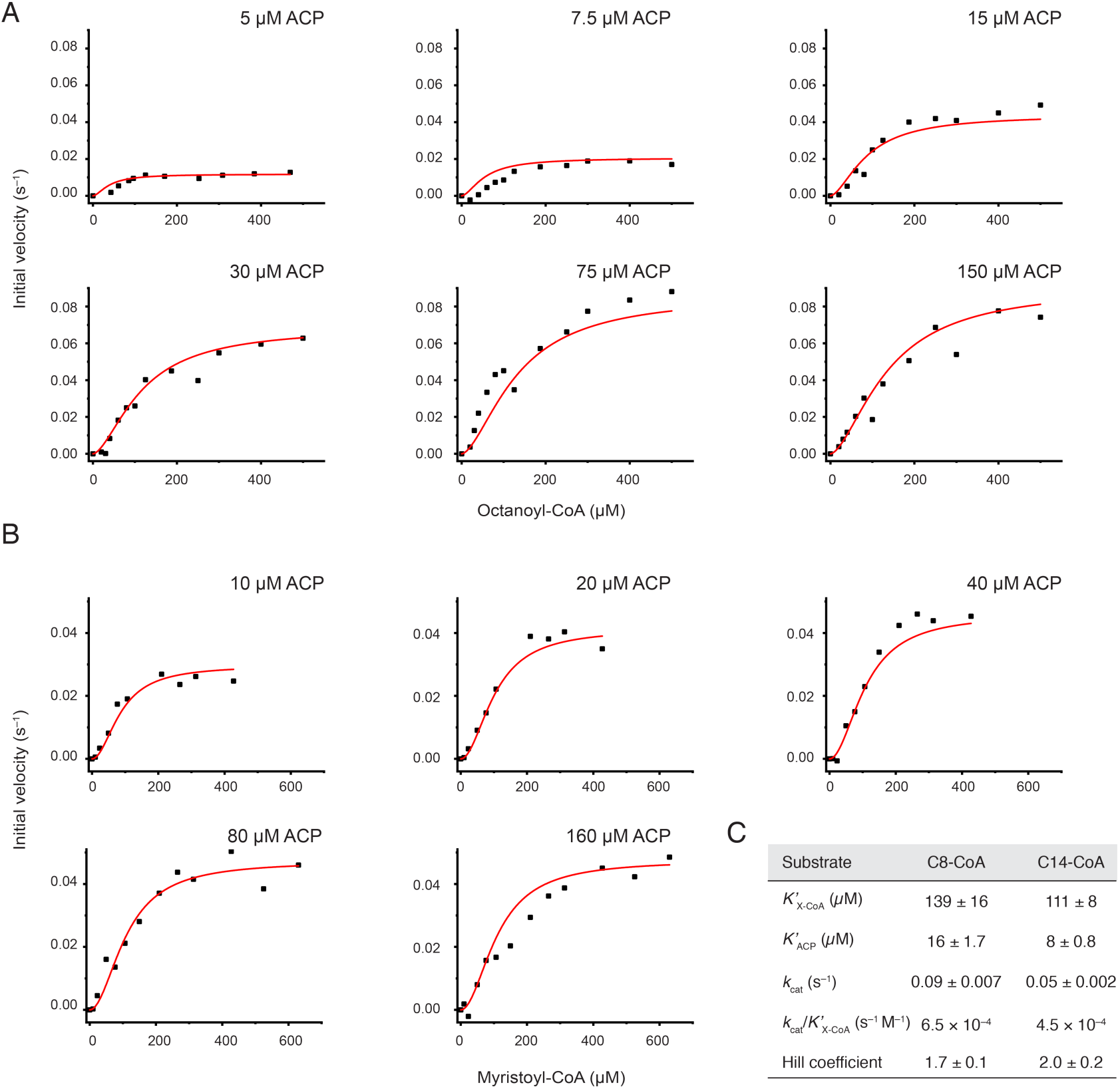
Comprehensive analysis of the KS-mediated transfer of octanoyl and myristoyl moieties. (A) Initial velocities plotted against octanoyl-CoA (C8-CoA) concentrations at six fixed ACP concentrations. (B) Initial velocities plotted against myristoyl-CoA (C14-CoA) concentrations at five fixed ACP concentrations. All data series were fit globally with the Hill equation due to the sigmoidal shape. (C) Absolute kinetic parameter derived from the respective global fits for octanoyl-CoA (C8-CoA) and myristoyl-CoA (C14-CoA), respectively. No parameters constraints were imposed during curve fitting. The constant *K*’ of the Hill equation is related to the Michaelis constant *K*_m_, but also contains terms related to the effect of substrate occupancy at one site on the substrate affinity of the other site.

## Discussion

We recently determined kinetic parameters for the murine MAT-mediated transfer of canonical and non-canonical acyl-CoA substrates illustrating the broad substrate tolerance of this domain (Rittner et al., 2018). How can this property be explained, considering high structural conservation to highly specific acyltransferases, like e. g. FabD of *E. coli*? The presented ensemble of MAT structures with non-covalently bound acyl-CoA and covalently bound acyl moieties shines light on this peculiarity. When considering the individual structures as snapshots of an overall conformational variability, data reveals significant dynamics within the MAT domain. Since soaking with octanoyl-CoA trapped the MAT in a very unusual conformation, revealing significant alterations in the position of active site residues, the newly presented structure is particularly informative in this respect. The data shows that substrate polyspecificity of MAT originates from the overall high relative positional dynamics of the α/β-hydrolase and the ferredoxin-like subdomain. A pronounced conformationally variability of the subdomain linkers SDL1 and SDL2, embedded in this large-scale, movement, is further relevant (Figure 3) for the accommodation of the chemically and structurally diverse substrates. Finally, residue R606 modulates substrate polyspecificity by either swinging out to liberate space for the acyl chain (e.g. octanoyl moieties), or by coordinating to the free carboxylic group of extender substrates (e.g. malonyl moieties) (Figure S6). This structural interpretation is supported by enzyme kinetic data as the (wildtype) murine MAT domain shows higher substrate ambiguity as the R606A-mutated MAT domain. In fact, the wildtype MAT domain is even able to accept octanoyl moieties with significantly higher efficiency than the R606A construct (Bunkoczi et al., 2009; Rittner et al., 2018), which could possibly be explained by a smaller number of populated conformational enzyme states (Table 2) (Khersonsky and Tawfik, 2010).

Can the rather strict KS domain substrate specificity be observed in structural properties? Indeed, the KS domain shows minor structural changes upon binding of an octanoyl chain, resembling a key-and-lock type binding, possessing strict specificities, as also observed for type II systems (von Wettstein-Knowles et al., 2006). The most prominent conformational change upon acylation with saturated acyl chains emerges from the stretch of residues 393-397, in particular residue F395, which is consequently slightly shifted in position and rotated in the side chain by approximately 125°. Its postulated role as a “gatekeeper” seems to be confirmed in animal FAS, as functional groups in the β-position of a bound acyl chain would sterically clash with the phenyl ring (Lee and Engels, 2014; Luckner et al., 2009). In the evolutionarily strongly related protein class of polyketide synthases (PKSs), which share a common KS domain fold, residues at the F395-equivalent position vary, reflecting the key feature of PKSs of condensing β-keto-, β-hydroxy- and α-β-unsaturated acyl substrates (Nguyen et al., 2008).

Our setup using a continuous coupled enzyme assay offers a convenient way to investigate the specificity of the KS domain in-depth. The kinetic characterization generally revealed maximum transacylation rates for CoA-esters of medium chain lengths (C8 to C12) and confirmed earlier results of Witkowski *et al*. (Figure 5B) (Witkowski et al., 1997). The unexpected low specific activity for short acyl chains may be attributed to the usage of CoA-esters as donors and substrate concentrations that are insufficient to fully saturate the enzyme. Titration of acyl-CoA substrates at different fixed ACP concentrations for octanoyl- and myristoyl-CoA resulted in sigmoidal individual initial velocity curves (Figure 6). Global fitting of the individual curves was possible when including Hill coefficients for both substrates and the obtained Hill-coefficients of 1.7-2 indicate positive cooperativity. This data can generally be interpreted by an increase in the efficiency of transferring an acyl chain to the active site cysteine of the KS domain, when the other KS of the dimer is already occupied. Such cooperativity of the domains of a KS dimer was postulated for the type II homologs (von Wettstein-Knowles et al., 2006) and could be explained by a conformational interconnection of both active sites that are pointing towards each other and merging at the two-fold axis.

Whereas a direct interaction of bound substrates can be ruled out as origin for cooperativity, because the enzyme bound acyl intermediates are too far apart from each other (15.6 Å between terminal C-atoms of octanoyl chains), a stretch of residues (residues 393-397) harboring the gatekeeping F395 and shifting upon acylation may be responsible for this phenomenon. The residues are residing at the dimeric interface and interact with a helix-turn-helix motif of the adjacent protomer (Figure 7A). In the acylated state, side chains of residues M132, Q136 and M139 (adjacent protomer) are slightly altered in their positions due to the rotation of F395, which furthermore leads to a slight shift of the turn (of the helix-turn-helix motif) by up to 0.7 Å (Figure 7B). Structural data reveals a putative coupling of this local rearrangement with active site residues via two hydrogen bonds; one between the side chains of R137 and D158 and the other between the carboxy group of D158 and the backbone amide of A160 (Figure 7C). All three residues are fully conserved in FASs. Based on this structural analysis, cooperativity could hence originate from a subtle reorganization of the active site residues in the neighboring protomer essentially induced by structural changes at the dimer interface occurring during acylation. Further experiments need to elaborate the molecular basis for the observed cooperative behavior of the KS domain as well as to analyze whether cooperativity observed for the transacylation step also extends to the Claisen condensation step.

**Figure 7:**
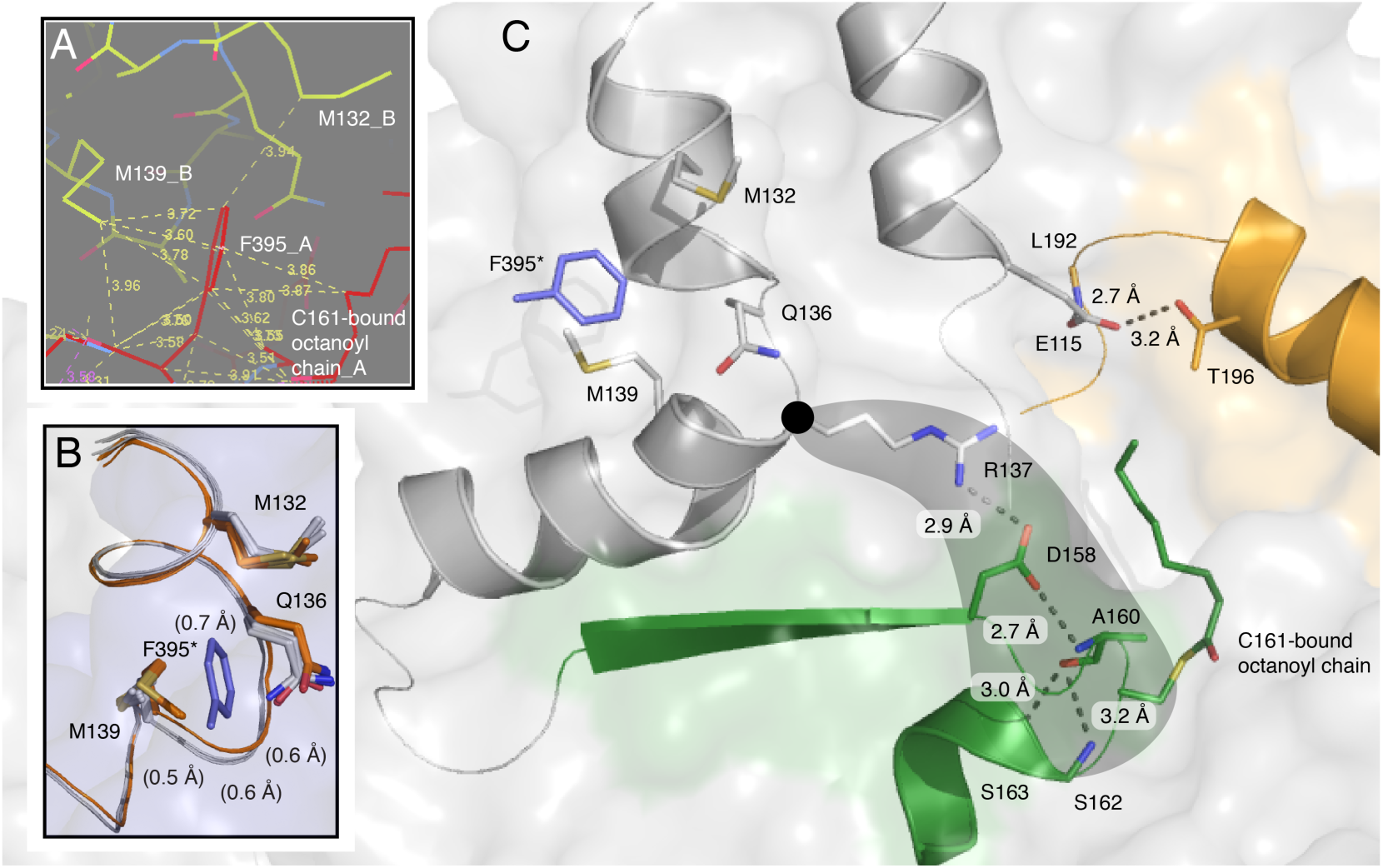
Structural interconnection of both KS active sites of a dimer. (A) Identification of M132 and M139 as F395-interacting residues in the other chain of the dimer. Chain A and chain B are colored in red and yellow, respectively. Analysis was performed with the software Coot. (B) Different side-chain conformations of M132, Q136 and M139 and subtle backbone shift between unbound (orange) and octanoyl-bound (grey) KS active sites. All four chains of both crystal structures (5my0 and 6rop) were aligned by a KS based superposition (BB of residues 1–407). F395 (blue) is depicted for clarity and the asterisks indicates the hypothetical position of the residue in the respective other protomer. (C) Hydrogen bonding between R137, D158 and the nucleophilic elbow (green) including the active site residue C161.

The specific kinetic information about the MAT and KS domains allow us to vividly draw the murine FAS function *in vivo*. The direct loading of the KS domain with acyl-CoA is not relevant *in vivo*, because the specificity constants of the KS-mediated transfer of acyl moieties from acyl-CoAs is more than three orders of magnitude lower than of the respective MAT-mediated transfer (Rittner et al., 2018). Accordingly, substrates are in general loaded at ACP by MAT-mediated transacylation. ACP-bound acetyl- and butyryl moieties are then transferred relatively slowly from the ACP domain to the KS domain and can in a competing pathway escape from FAS by MAT-mediated offloading. Release of short acyl-CoAs via the MAT domain depends on the *in vivo* ratio between malonyl-CoA, acetyl-CoA and free coenzyme A and hence the availability of an empty MAT’s active site. With increasing malonyl-CoA concentrations at higher energetic state of the cell, the offloading event of acetyl- and butyryl moieties gets more and more unlikely and further elongation becomes dominant (Abdinejad et al., 1981). As shown here, once a chain length of six carbons and longer is reached, transacylation from the ACP domain to the KS domain becomes highly efficient until it sharply drops with a chain length of 16 carbons (Figure 5B). According to a key finding of this study, the efficiency of FAS in the elongation of the acyl chain is increased when both reaction chambers are used simultaneously, as acylation of one KS domain of the FAS dimer accelerates the acylation of the other (Figure 6). The substrate specificity of the KS domain is important for the specific production of palmitic acid and is assisted by the substrate specificity of the TE domain for long fatty acyl intermediates (Cheng et al., 2008; Heil et al., 2019; Naggert et al., 1991; Zhang et al., 2011). In summary, FAS produces short acetyl and butyryl-CoA esters in a low energetic state of the cell and almost exclusively palmitic acid (C16) at high energetic states.

The presented structural and mechanistic study deepens our molecular understanding of the two initial catalytic domains in animal fatty acid synthesis. Such detailed information is particularly interesting as FAS got into focus as a target for combinatorial anti-cancer therapy. Especially, the plasticity of the MAT domain shall be highlighted, allowing to accommodate a broad range of chemically diverse compounds. This may aid future rational drug discovery campaigns and enlarge the pool of potentially screened lead structures.

## Acknowledgments

This work was supported by a Lichtenberg grant of the Volkswagen Foundation to M.G. (grant number 85701). Further support was received from the LOEWE program (Landes-Offensive zur Entwicklung wissenschaftlich-ökonomischer Exzellenz) of the state of Hesse conducted within the framework of the MegaSyn Research Cluster.

## Authors contribution

A.R. performed protein expression, purification experiments, enzymatic assays and analyzed corresponding data. A.R. conceived the project. M.G. designed the research. A.H. performed kinetic experiments with the KS domain using octanoyl-CoA under supervision of A.R. Crystallization was performed by A.R., A.H. and K.S.P. Crystal structure was solved by K.S.P. A.R., K.S.P. and M.G. analyzed data and wrote the manuscript.

## Declaration of interests

The authors declare no competing interests.

## STAR Methods

### Key Resources Table

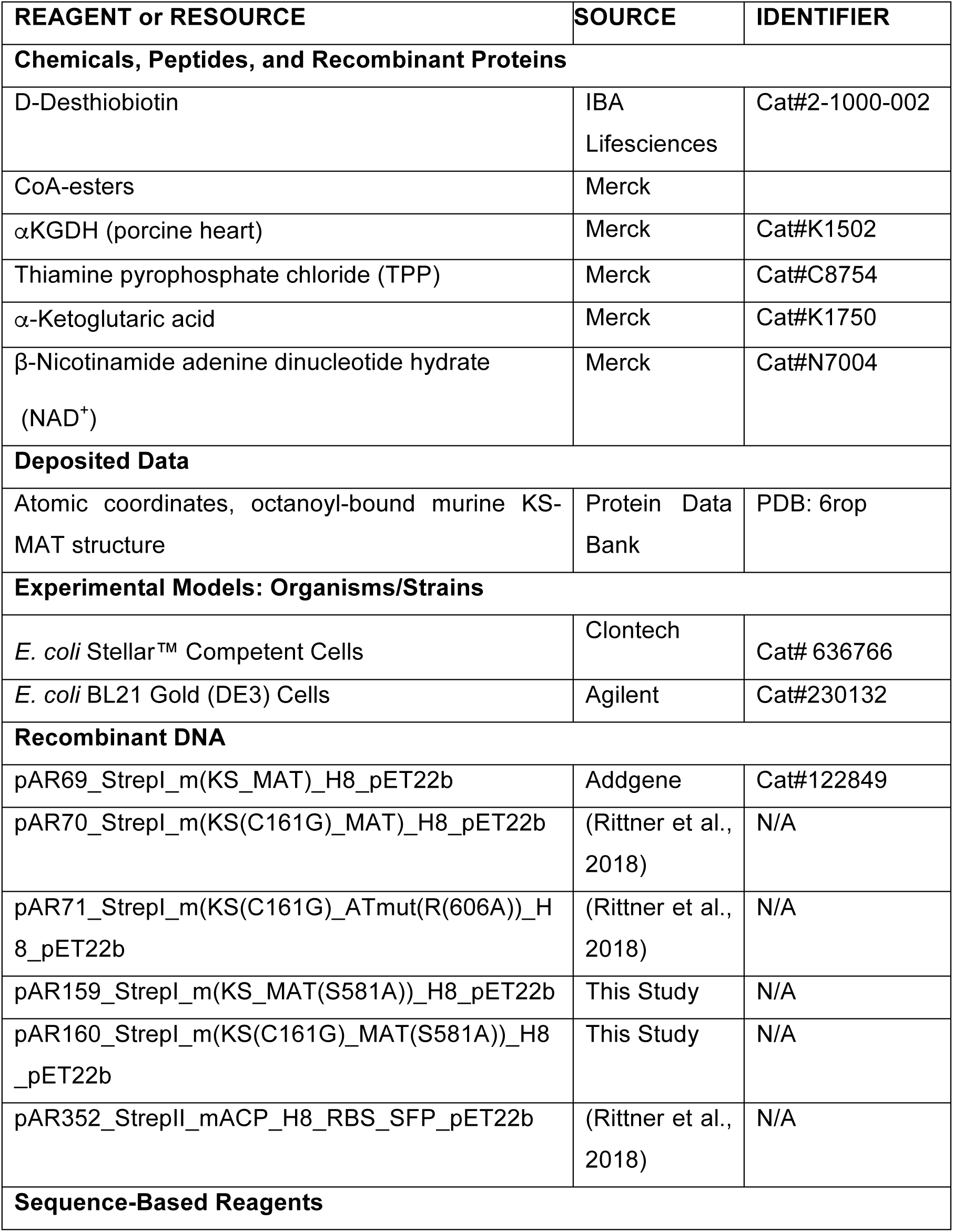
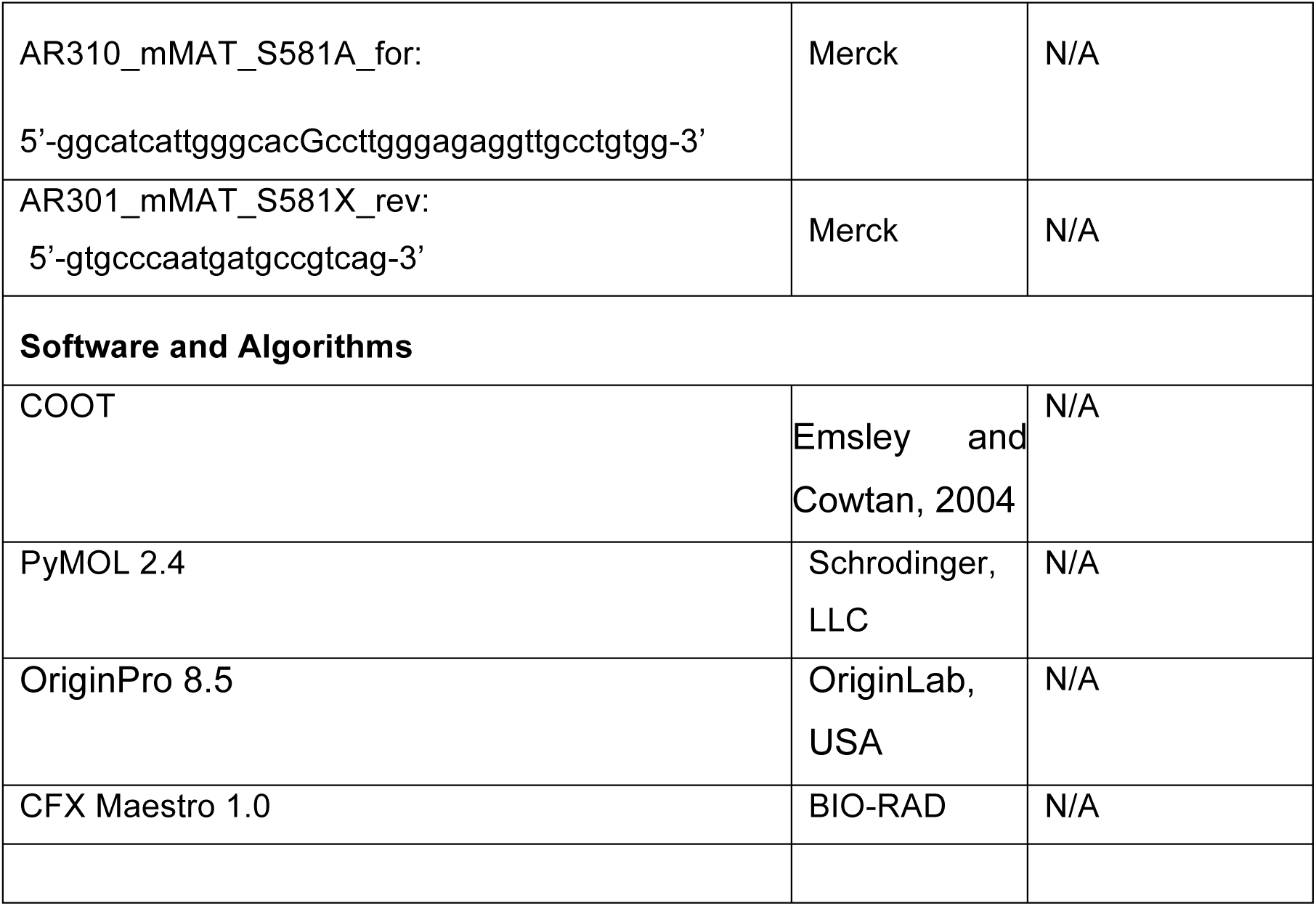

### Contact for Reagent and Resource Sharing

Further information and requests for reagents may be directed to, and will be fulfilled by the corresponding author Martin Grininger.

### Experimental Model and Subject Details

#### Method Details

##### Reagents and Constructs

All CoA-esters, β -NAD^+^, NADH, α -ketoglutarate dehydrogenase (porcine heart) (αKGDH), α-ketoglutaric acid, thiamine pyrophosphate (TPP), and EDTA were purchased from Merck. BSA was from Serva. Restriction enzymes were bought from NEB biolabs. IPTG was from Carl Roth. Ni-NTA affinity resin was from Qiagen and 5 mL Strep-Tactin→ columns were purchased from IBA technologies. Purity of CoA-esters was confirmed by HPLC-UV analysis before usage.

Point mutations were introduced by PCR based cloning. Fragments for pAR159 and pAR160 were generated by amplification of pAR69 (Addgene, #122849) and pAR70 (Rittner et al., 2018) with the primer pair: AR301 (5’-gtgcccaatgatgccgtcag-3’) and AR310 (5’-ggcatcattgggcacGccttgggagaggttgcctgtgg-3’). PCR products were treated with Dpn1 (NEB), purified by gel electrophoresis and DNA was extracted with the Wizard® SV Gel and PCR Clean-Up System (Promega). Purified DNA was transformed into *E. coli* Stellar™ Competent Cells, 5 mL LB cultures were grown and plasmids were isolated with the PureYield™ Plasmid Miniprep System (Promega). Sequences of all plasmids were confirmed with the “dye terminator” method.

##### Expression and purification of KS-MAT variants

All plasmids were transformed into chemically competent *E. coli* BL21 Gold (DE3) cells. The transformants were grown overnight at 37 °C in 20 mL LB (100 µg mL^−1^ ampicillin (amp) and 1 % (w/v) glucose) medium. Pre-cultures were used to inoculate 1 L TB medium (100 µg mL^−1^ amp). Cultures were grown at 37 °C until they reached an optical density (OD_600_) of 0.5–0.6. After cooling at 4 °C for 20 min, cultures were induced with 0.25 mM IPTG, and grown for additional 16 h at 20 °C and 180 rpm. Cells were harvested by centrifugation (4,000 rcf for 20 min). The cell pellets were resuspended in lysis buffer (50 mM potassium phosphate, 200 mM potassium chloride, 10 % (v/v) glycerol, 1 mM EDTA, 30 mM imidazole (pH 7.0)) and lysed by French press. After centrifugation at 50,000 rcf for 30 min, the supernatant was mixed with 1 M MgCl_2_ to a final concentration of 2 mM. The cytosol was transferred to Ni-NTA-columns and washed with 5 column volumes (CV) wash buffer (lysis buffer without EDTA). Bound protein was eluted with 2.5 CV elution buffer (50 mM potassium phosphate, 200 mM potassium chloride, 10 % (v/v) glycerol, 300 mM imidazole (pH 7.0). The eluent was transferred to Strep-Tactin-columns, and washed with 5 CV strep-wash buffer (250 mM potassium phosphate, 10 % (v/v) glycerol, 1 mM EDTA, (pH 7.0)). Proteins were eluted with 2.5 CV elution buffer (strep-wash buffer containing 2.5 mM D-desthiobiotin). After concentration to 10–20 mg mL^−1^, the proteins were frozen in liquid nitrogen and stored at –80 °C. Samples were thawn at 37 °C for 30 min and further polished by size-exclusion chromatography (SEC) using a Superdex 200 GL10/300 column equilibrated with the strep-wash buffer. Proteins were concentrated again to 10–20 mg mL^−1^ and stored frozen in aliquots using liquid nitrogen.

##### Expression and purification of ACP

ACP for the activity assay was produced by co-expression with the 4’-phosphopantetheinyl transferase Sfp from the bicistronically organized vector pAR352 (Rittner et al., 2018).The plasmid was transformed into chemically competent *E. coli* BL21 Gold (DE3) cells. Overnight cultures were grown in 40 mL LB (100 µg mL^−1^ ampicillin (amp) and 1 % (w/v) glucose) at 37 °C. 2 L TB medium (100 µg mL^−1^ amp) was inoculated with the overnight culture and incubated at 37 °C until an optical cell density (OD_600_) of 0.5–0.6 was reached. After cooling at 4 °C for 20 min, cultures were induced with 0.25 mM IPTG, and grown for additional 16 h at 20 °C and 180 rpm. Cells were harvested by centrifugation, resuspended in lysis buffer and lysed by French press. After centrifugation (50,000 rcf for 30 min), the supernatant (supplemented with 2 mM MgCl_2_) was transferred to Ni-NTA-columns and washed with 5 CV wash buffer. The protein was eluted with elution buffer (wash buffer containing 300 mM imidazole) and concentrated. Pooled fractions, were separated on a Superdex 200 HiLoad 16/60 or 26/60 SEC column equilibrated with buffer (50 mM potassium phosphate, 200 mM potassium chloride, 10 % (v/v) glycerol, 1 mM EDTA). All fractions containing monomeric ACP were pooled and concentrated to 10-20 mg mL^−1^.

#### Protein concentration

Protein concentrations were calculated from the absorbance at 280 nm, which was recorded on a NanoDrop 2000c (Thermo scientific). Extinction coefficients were calculated from the primary sequence without *N*-formylmethionine with CLC Main workbench (Qiagen). Absorbance 1 g L^−1^ at 280 nm (10 mm): 1.053 for KS-MAT and 0.475 for ACP.

##### Crystallization, data collection and structure determination

Crystallization conditions for KS-MAT were as previously published (Rittner et al., 2018). Single-crystals were obtained at 0.2 M potassium-sodium tartrate, 25 % (w/v) PEG 3350 at 20 °C to sizes of about 75 × 75 × 75 µm^3^. Drops with the crystals were supplemented with 0.5 µl of 10 mM octanoyl-CoA (Merck) for up to 2 minutes and subsequently treated with a cryosolution containing 20 % (v/v) glycerol in the mother liquor. The crystal was then picked in a nylon fiber loop and vitrified into liquid nitrogen. Single wavelength X-radiation diffraction dataset was collected at the Swiss Light Source (X06SA), and maintained at 100 K, while data were recorded onto a detector (DECTRIS EIGER 16M). Using the ‘goeiger.com’ pipeline at X06SA, data reduction was performed within *XDS* (Kabsch, 2010), for indexing and integration, and *Aimless* (Evans, 2005) for scaling. The structural model of a monomer from the murine FAS KS-MAT didomain complexed with Malonyl-CoA (pdb accession code 5my0) was used to solve the phase problem using the program *Molrep* (Vagin and Teplyakov, 1997). After an initial rigid-body refinement, the model was subjected to repeated cycles of restrained refinement with *REFMAC*5 (Murshudov et al., 2011) with manual model building using *Coot* (Emsley et al., 2010). Data collection and refinement statistics are given in Table 1. Electron density maps were generated by Phenix (Afonine et al., 2012).

##### Thermal shift assay

Thermal shift assays were performed as previously reported (Rittner et al., 2019). Briefly, 2 µL of protein solution (20 µM) were mixed with 21 µL of the respective buffer and 2 µL of SYPRO Orange protein gel stain (5000 × diluted), then fluorescence was measured from 5 °C to 95 °C with a step of 0.5 °C min^−1^, with excitation wavelength set to 450-490 nm, and emission wavelength to 560-580 nm. Data was analyzed with the software CFX Maestro 1.0.

##### α-Ketoglutarate dehydrogenase coupled activity assay

The enzyme-coupled assay was performed as previously published (Rittner et al., 2018), which was adapted from reference (Molnos et al., 2003). Assays (octanoyl-CoA transacylation of the KS^C161G^-MAT^R606A^ variant and the series of response curves for the KS-mediated transacylation of octanoyl moieties) were run in 96-well f-bottom microtiter plates (Greiner Bio-one) and NADH fluorescence was monitored using a ClarioStar microplate reader equipped with a dispenser (BMG labtech) at the following settings; excitation: 348-20 nm; emission: 476-20 nm; gain: 1900; focal height: 5.2 mm; flashes: 70; orbital averaging: 4 mm.

Within this study we reduced the assay volume first to 50 µL in 96-well Half Area Microplates (Greiner Bio-one) and then to 20 µL in 384-well Small Volume HiBase Microplates (Greiner Bio-one) for practical and financial reasons. The chain-length specificity of the KS domain (Figure 5) and the series of response curves for the KS-mediated transacylation of myristoyl moieties (Figure 6B) were measured in half-area plates and 384-well plates, respectively. New calibration curves with NADH and control measurements were performed for both smaller plate formats. Settings for the microplate reader were for half-area plates: 348-20 nm; emission: 476-20 nm; gain: 1500; focal height: 5.6 mm; flashes: 17; orbital averaging: 1 mm; and for 384-well plates: 348-20 nm; emission: 476-20 nm; gain: 1500; focal height: 11.9 mm; flashes: 17; orbital averaging: off.

Same procedures were used in preparation of assays even if they had different volumes. Briefly, four different solutions were prepared in assay buffer (50 mM sodium phosphate, 10 % (v/v) glycerol, 1 mM DTT, 1 mM EDTA (pH 7.6), filtered and degased). Solution 1 (Sol 1) contained murine KS-MAT in a 3.33-fold or 4-fold concentrated stock solution and supplemented with 0.1 mg mL^−1^ BSA. Solution 2 (Sol 2) contained 8 mM α-ketoglutaric acid, 1.6 mM NAD^+^, 1.6 mM TPP and 60 mU/100 µL αKGDH, representing a 4-fold concentrated stock. Solution 3 (Sol 3) contained 4-fold concentrated CoA-esters, typically between 0.4–2800 µM. Solution 4 (Sol 4) finally contained 5-fold or 4-fold concentrated murine ACP, typically between 20-800 µM. The components were pipetted in order: 30 µL Sol 1 (3.33-fold), 25 µL Sol 2 and 25 µL Sol 3 for 96-well plates; 15 µL Sol 1 (3.33-fold), 12.5 µL Sol 2 and 12.5 µL Sol 3 for 96-well half-area plates and 5 µL Sol 1 (4-fold), 5 µL Sol 2 and 5 µL Sol 3 for 384-well plates, followed by mixing. The transfer reaction was initiated by 20 µL and 10 µL Sol 4 (5-fold)) or 5 µL Sol 4 (4-fold), which was added by the dispenser. The final concentrations of all ingredients were 50 mM sodium phosphate, pH 7.6, 10 % (v/v) glycerol, 1 mM DTT, 1 mM EDTA, 2 mM α-ketoglutaric acid, 0.4 mM NAD^+^, 0.4 mM TPP, 15 mU/100 µL αKGDH, 0.03 mg mL^−1^ BSA, 100-200 nM KS-MAT, 5–160 µM ACP, 0.1–700 µM X-CoA (where X refers to the respective acyl-moiety of the assay). The background noise of the assay set-up was determined with assay buffer supplemented with 0.1 mg mL^−1^ BSA. Equidistant kinetic measurements were taken every 5-22 s for ca. 5 min at 30 °C.

##### Transacylation kinetics of the MAT^R606A^ variant for octanoyl moieties

Determining the apparent Michaelis-Menten constant is an iterative process. Pre-experiments were initially performed to approach the approximate value of *K*_m_. Final concentrations of enzyme and ACP were 100 nM and 60 µM, respectively. Eight data points were collected that cover substrate concentrations (Sol 3) of 0.2 × *K*_m_; 0.3 × *K*_m_; 0.5 × *K*_m_; 0.75 × *K*_m_; 1.25 × *K*_m_; 2 × *K*_m_; 3 × *K*_m_; 5 × *K*_m_. Every measurement was performed in technical triplicates and the corresponding background signal was substracted. Experiments were setup in a way such that changes in signal remained linear during the time ranges of measurement. Data were collected in biological replicates (n = 4) and were fit with the Michaelis-Menten function using OriginPro 8.5 (OriginLab, USA).

##### Chain-length specificity of the KS domain

KS-mediated transacylation of acyl-CoAs were measured at fixed enzyme (200 nM), ACP (75 µM) and acyl-CoA (500 µM) concentrations. Turnover rates were determined by linear fit and error bars reflect the standard deviation from three biological replicates (n = 3).

##### Analysis of KS-mediated transfer of octanoyl and myristoyl moieties

Initial velocities were determined for twelve different CoA-ester concentrations at six (octanoyl-CoA) and five (myristoyl-CoA) fixed ACP concentrations. Enzyme concentration was 200 nM. Every measurement was performed in one biological replicate (n = 1) and the corresponding background signal was subtracted. Both series of response curves were globally fit using all data without any parameter constraints. The global fit was performed with OriginPro 8.5 (OriginLab, USA) using the following equations for the ping-pong mechanism:

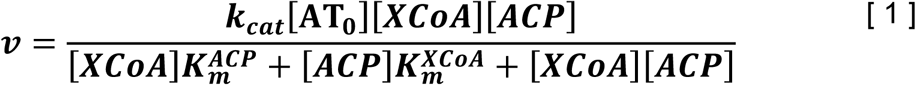

Hill-type variation:

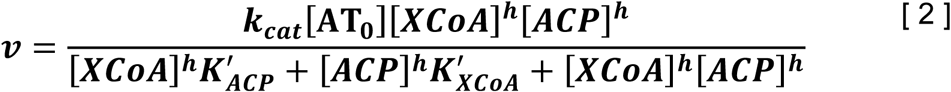

### Quantification and Statistical Analysis

Statistical parameters are reported in Figure Legends and in Method Details. Biological replicates refer to independently expressed and purified enzymes. Transacylation kinetics of the MAT^R606A^ variant for octanoyl moieties was performed in four biological replicates, with error bars indicating SD. Chain-length specificity of the KS domain was performed in three biological replicates, with error bars indicating SD. Both series of response curves were performed in one biological replicate and the errors are standard errors from the global fit.

### Data and Software Availability

Accession numbers for the atomic coordinates for the octanoyl-bound murine KS-MAT structure reported in this paper is PDB: 6rop. X-ray diffraction data are publicly available at https://zenodo.org/deposit/2785017.

### Supplementary Material

Additional supporting information may be found in the online version of this article.

## References

Abdinejad, A., Fisher, A.M., and Kumar, S. (1981). Production and Utilization of Butyryl-CoA by Fatty Acid Synthetase from Mammalian Tissues. Arch. Biochem. Biophys. 208, 135–145.

Afonine, P.V., Grosse-Kunstleve, R.W., Echols, N., Headd, J.J., Moriarty, N.W., Mustyakimov, M., Terwilliger, T.C., Urzhumtsev, A., Zwart, P.H., and Adams, P.D. (2012). Towards automated crystallographic structure refinement with phenix.refine. Acta Cryst (2012). D68, 352–367 [doi:10.1107/S0907444912001308], 1-16.

Afonine, P.V., Moriarty, N.W., Mustyakimov, M., Sobolev, O.V., Terwilliger, T.C., Turk, D., Urzhumtsev, A., and Adams, P.D. (2015). FEM: feature-enhanced map. Acta Crystallogr. D Biol. Crystallogr. 71, 646–666.

Beld, J., Lee, D.J., and Burkart, M.D. (2015). Fatty acid biosynthesis revisited: structure elucidation and metabolic engineering. Mol. Biosyst. 11, 38–59.

Boehringer, D., Ban, N., and Leibundgut, M. (2013). 7.5-Å Cryo-EM Structure of the Mycobacterial Fatty Acid Synthase. J. Mol. Biol., 1-9.

Buckley, D., Duke, G., Heuer, T.S., O’Farrell, M., Wagman, A.S., McCulloch, W., and Kemble, G. (2017). Fatty acid synthase - Modern tumor cell biology insights into a classical oncology target. Pharmacol. Ther. 177, 23–31.

Buckner, J.S., Kolattukudy, P.E., and Rogers, L. (1978). Synthesis of Multimethyl-Branched Fatty Acids by Avian and Mammalian Fatty Acid Synthetase and Its Regulation by Malonyl-CoA Decarboxylase in the Uropygial Gland. Arch. Biochem. Biophys. 186, 152–163.

Bunkoczi, G., Misquitta, S., Wu, X., Lee, W.H., Rojkova, A., Kochan, G., Kavanagh, K.L., Oppermann, U., and Smith, S. (2009). Structural Basis for Different Specificities of Acyltransferases Associated with the Human Cytosolic and Mitochondrial Fatty Acid Synthases. Chem. Biol. 16, 667–675.

Chen, A., Re, R.N., and Burkart, M.D. (2018). Type II fatty acid and polyketide synthases: deciphering protein–protein and protein–substrate interactions. Nat. Prod. Rep. 35, 1029–1045.

Cheng, F., Wang, Q., Chen, M., Quiocho, F.A., and Ma, J. (2008). Molecular docking study of the interactions between the thioesterase domain of human fatty acid synthase and its ligands. Proteins: Structure, Function, and Bioinformatics 70, 1228–1234.

Copeland, R.A. (2005). Enzymes - A Practical Introduction to Structure, Mechanism and Data Analysis - 2nd Edition (Weinheim, Germany: Wiley-VCH Verlag GmbH).

Dean, E.J., Falchook, G.S., Patel, M.R., Brenner, A.J., Infante, J.R., Arkenau, H.-T., Borazanci, E.H., Lopez, J.S., Pant, S., Schmid, P., et al. (2016). Preliminary activity in the first in human study of the first-in-class fatty acid synthase (FASN) inhibitor, TVB-2640. J. Clin. Oncol. 34, 2512–2512.

Dodge, G.J., Patel, A., Jaremko, K.L., McCammon, J.A., Smith, J.L., and Burkart, M.D. (2019). Structural and dynamical rationale for fatty acid unsaturation in *Escherichia coli*. Proc. Natl. Acad. Sci. USA 116, 6775–6783.

Elad, N., Baron, S., Peleg, Y., Albeck, S., Grunwald, J., Raviv, G., Shakked, Z., Zimhony, O., and Diskin, R. (2018). Structure of Type-I Mycobacterium tuberculosis fatty acid synthase at 3.3 Å resolution. Nature Communications 9, 3886.

Emsley, P., Lohkamp, B., Scott, W.G., and Cowtan, K. (2010). Features and development of *Coot*. Acta Crystallogr. D Biol. Crystallogr. 66, 486–501.

Evans, P. (2005). Scaling and assessment of data quality. Acta Crystallogr. D Biol. Crystallogr. 62, 72–82.

Flocco, M.M., and Mowbray, S.L. (1994). Planar stacking interactions of arginine and aromatic side-chains in proteins. J. Mol. Biol. 235, 709–717.

Gansler, T.S., Hardman, W., Hunt, D.A., Schaffel, S., and Hennigar, R.A. (1997). Increased expression of fatty acid synthase (OA-519) in ovarian neoplasms predicts shorter survival. Hum. Pathol. 28, 686–692.

Grininger, M. (2014). Perspectives on the evolution, assembly and conformational dynamics of fatty acid synthase type I (FAS I) systems. Curr. Opin. Struct. Biol. 25, 49–56.

Hardwicke, M.A., Rendina, A.R., Williams, S.P., Moore, M.L., Wang, L., Krueger, J.A., Plant, R.N., Totoritis, R.D., Zhang, G., Briand, J., et al. (2014). A human fatty acid synthase inhibitor binds β-ketoacyl reductase in the keto-substrate site. Nat. Chem. Biol. 10, 774–779.

Heil, C.S., Wehrheim, S.S., Paithankar, K.S., and Grininger, M. (2019). Fatty acid biosynthesis: Chain length regulation and control. ChemBioChem 0.

Herbst, D.A., Townsend, C.A., and Maier, T. (2018). The architectures of iterative type I PKS and FAS. Nat. Prod. Rep. 35, 1046–1069.

Hol, W.G. (1985). The role of the alpha-helix dipole in protein function and structure. Prog. Biophys. Mol. Biol. 45, 149–195.

Johansson, P., Wiltschi, B., Kumari, P., Kessler, B., Vonrhein, C., Vonck, J., Oesterhelt, D., and Grininger, M. (2008). Inhibition of the fungal fatty acid synthase type I multienzyme complex. Proceedings of the National Academy of Sciences 105, 12803–12808.

Kabsch, W. (2010). XDS. Acta Crystallogr. D Biol. Crystallogr. 66, 125–132

Khandekar, M.J., Cohen, P., and Spiegelman, B.M. (2011). Molecular mechanisms of cancer development in obesity. Nature Reviews Cancer 11, 886.

Khersonsky, O., and Tawfik, D.S. (2010). Enzyme Promiscuity: A Mechanistic and Evolutionary Perspective. Annu. Rev. Biochem. 79, 471–505.

Kuhajda, F.P. (2006). Fatty Acid Synthase and Cancer: New Application of an Old Pathway. Cancer Res. 66, 5977–5980.

Lee, W., and Engels, B. (2014). The Protonation State of Catalytic Residues in the Resting State of KasA Revisited: Detailed Mechanism for the Activation of KasA by Its Own Substrate. Biochemistry 53, 919–931.

Leibundgut, M., Jenni, S., Frick, C., and Ban, N. (2007). Structural Basis for Substrate Delivery by Acyl Carrier Protein in the Yeast Fatty Acid Synthase. Science 316, 288–290.

Liebschner, D., Afonine, P.V., Moriarty, N.W., Poon, B.K., Sobolev, O.V., Terwilliger, T.C., and Adams, P.D. (2017). Polder maps: improving OMIT maps by excluding bulk solvent. Acta Cryst (2017). D73, 148–157 [doi:10.1107/S2059798316018210], 1-10.

Lomakin, I.B., Xiong, Y., and Steitz, T.A. (2007). The Crystal Structure of Yeast Fatty Acid Synthase, a Cellular Machine with Eight Active Sites Working Together. Cell 129, 319–332.

Luckner, S.R., Machutta, C.A., Tonge, P.J., and Kisker, C. (2009). Crystal Structures of *Mycobacterium tuberculosis* KasA Show Mode of Action within Cell Wall Biosynthesis and its Inhibition by Thiolactomycin. Structure/Folding and Design 17, 1004–1013.

Maier, T., Leibundgut, M., and Ban, N. (2008). The Crystal Structure of a Mammalian Fatty Acid Synthase. Science 321, 1315–1322.

Maier, T., Leibundgut, M., Boehringer, D., and Ban, N. (2010). Structure and function of eukaryotic fatty acid synthases. Q. Rev. Biophys. 43, 373–422.

Menendez, J.A., Vazquez-Martin, A., Ortega, F.J., and Fernandez-Real, J.M. (2009). Fatty acid synthase: association with insulin resistance, type 2 diabetes, and cancer. Clin. Chem. 55, 425–438.

Molnos, J., Gardiner, R., Dale, G.E., and Lange, R. (2003). A continuous coupled enzyme assay for bacterial malonyl–CoA:acyl carrier protein transacylase (FabD). Anal. Biochem. 319, 171–176.

Murshudov, G.N., Skubák, P., Lebedev, A.A., Pannu, N.S., Steiner, R.A., Nicholls, R.A., Winn, M.D., Long, F., and Vagin, A.A. (2011). *REFMAC5* for the refinement of macromolecular crystal structures. Acta Crystallogr. D Biol. Crystallogr. 67, 355–367.

Naggert, J., Witkowski, A., Wessa, B., and Smith, S. (1991). Expression in Escherichia coli, purification and characterization of two mammalian thioesterases involved in fatty acid synthesis. Biochem. J. 273, 787–790.

Nguyen, P.L., Ma, J., Chavarro, J.E., Freedman, M.L., Lis, R., Fedele, G., Fiore, C., Qiu, W., Fiorentino, M., Finn, S., et al. (2010). Fatty Acid Synthase Polymorphisms, Tumor Expression, Body Mass Index, Prostate Cancer Risk, and Survival. J. Clin. Oncol. 28, 3958–3964.

Nguyen, T., Ishida, K., Jenke-Kodama, H., Dittmann, E., Gurgui, C., Hochmuth, T., Taudien, S., Platzer, M., Hertweck, C., and Piel, J. (2008). Exploiting the mosaic structure of *trans*-acyltransferase polyketide synthases for natural product discovery and pathway dissection. Nat. Biotechnol. 26, 225–233.

Olsen, J.G., Kadziola, A., von Wettstein-Knowles, P., Siggaard-Andersen, M., and Larsen, S. (2001). Structures of beta-ketoacyl-acyl carrier protein synthase I complexed with fatty acids elucidate its catalytic machinery. Structure/Folding and Design 9, 233–243.

Paiva, P., Sousa, S.r.F., Ramos, M.J., and Fernandes, P.A. (2018). Understanding the Catalytic Machinery and the Reaction Pathway of the Malonyl-Acetyl Transferase Domain of Human Fatty Acid Synthase. ACS Catalysis 8, 4860–4872.

Pandey, P.R., Liu, W., Xing, F., Fukuda, K., and Watabe, K. (2012). Ant-Cancer Drugs Targeting Fatty Acid Synthase (FAS). Recent Pat. Anticancer Drug Discov. 7, 185–197.

Pappenberger, G., Benz, J., Gsell, B., Hennig, M., Ruf, A., Stihle, M., Thoma, R., and Rudolph, M.G. (2010). Structure of the Human Fatty Acid Synthase KS–MAT Didomain as a Framework for Inhibitor Design. J. Mol. Biol. 397, 508–519.

Pemble, C.W., Johnson, L.C., Kridel, S.J., and Lowther, W.T. (2007). Crystal structure of the thioesterase domain of human fatty acid synthase inhibited by Orlistat. Nature Structural &#38; Molecular Biology 14, 704–709.

Ploskoń, E., Arthur, C.J., Evans, S.E., Williams, C., Crosby, J., Simpson, T.J., and Crump, M.P. (2008). A Mammalian Type I Fatty Acid Synthase Acyl Carrier Protein Domain Does Not Sequester Acyl Chains. J. Biol. Chem. 283, 518–528.

Rangan, V.S., and Smith, S. (1997). Alteration of the Substrate Specificity of the Malonyl-CoA/Acetyl-CoA: Acyl Carrier Protein *S*-Acyltransferase Domain of the Multifunctional Fatty Acid Synthase by Mutation of a Single Arginine Residue. J. Biol. Chem. 272, 11975–11978.

Rashid, A., Pizer, E.S., Moga, M., Milgraum, L.Z., Zahurak, M., Pasternack, G.R., Kuhajda, F.P., and Hamilton, S.R. (1997). Elevated expression of fatty acid synthase and fatty acid synthetic activity in colorectal neoplasia. The American journal of pathology 150, 201–208.

Rittner, A., Paithankar, K.S., Drexler, D.J., Himmler, A., and Grininger, M. (2019). Probing the modularity of megasynthases by rational engineering of a fatty acid synthase Type I. Protein Sci. 28, 414–428.

Rittner, A., Paithankar, K.S., Huu, K.V., and Grininger, M. (2018). Characterization of the Polyspecific Transferase of Murine Type I Fatty Acid Synthase (FAS) and Implications for Polyketide Synthase (PKS) Engineering. ACS Chem. Biol. 13, 723–732.

Rossini, E., Gajewski, J., Klaus, M., Hummer, G., and Grininger, M. (2018). Analysis and engineering of substrate shuttling by the acyl carrier protein (ACP) in fatty acid synthases (FASs). Chem. Commun. 68, 501.

Semenkovich, C.F., Coleman, T., and Fiedorek, F.T. (1995). Human fatty acid synthase mRNA: tissue distribution, genetic mapping, and kinetics of decay after glucose deprivation. J. Lipid Res. 36, 1507–1521.

Smith, S., and Stern, A. (1983). The Effect of Aromatic CoA Esters on Fatty Acid Synthetase: Biosynthesis of *ω*-Phenyl Fatty Acids. Arch. Biochem. Biophys. 222, 259–265.

Smith, S., and Tsai, S.-C. (2007). The type I fatty acid and polyketide synthases: a tale of two megasynthases. Nat. Prod. Rep. 24, 1041–1072.

Sztain, T., Bartholow, T.G., Orcid, J.A.M., and Orcid, M.D.B. (2019). Shifting the Hydrolysis Equilibrium of Substrate Loaded Acyl Carrier Proteins. Biochemistry, 1–4.

Uhlén, M., Fagerberg, L., Hallström, B.M., Lindskog, C., Oksvold, P., Mardinoglu, A., Sivertsson, Å., Kampf, C., Sjöstedt, E., Asplund, A., et al. (2015). Tissue-based map of the human proteome. Science 347, 1260419.

Vagin, A., and Teplyakov, A. (1997). *MOLREP*: an Automated Program for Molecular Replacement. J. Appl. Crystallogr. 30, 1022–1025.

von Wettstein-Knowles, P., Olsen, J.G., McGuire, K.A., and Henriksen, A. (2006). Fatty acid synthesis. Role of active site histidines and lysine in Cys-His-His-type beta-ketoacyl-acyl carrier protein synthases. FEBS J. 273, 695–710.

Wang, J., Soisson, S.M., Young, K., Shoop, W., Kodali, S., Galgoci, A., Painter, R., Parthasarathy, G., Tang, Y.S., Cummings, R., et al. (2006). Platensimycin is a selective FabF inhibitor with potent antibiotic properties. Nature 441, 358–361.

White, S.W., Zheng, J., Zhang, Y.-M., and Rock, C.O. (2005). The Structural Biology of Type II Fatty Acid Biosynthesis. Annu. Rev. Biochem. 74, 791–831.

Winn, M.D., Isupov, M.N., and Murshudov, G.N. (2001). Use of TLS parameters to model anisotropic displacements in macromolecular refinement. Acta Crystallographica Section D 57, 122–133.

Witkowski, A., Joshi, A.K., and Smith, S. (1997). Characterization of the Interthiol Acyltransferase Reaction Catalyzed by the *β*-Ketoacyl Synthase Domain of the Animal Fatty Acid Synthase. Biochemistry 36, 16338–16344.

Zhang, W., Chakravarty, B., Zheng, F., Gu, Z., Wu, H., Mao, J., Wakil, S.J., and Quiocho, F.A. (2011). Crystal structure of FAS thioesterase domain with polyunsaturated fatty acyl adduct and inhibition by dihomo-γ-linolenic acid. Proc. Natl. Acad. Sci. U. S. A. 108, 15757–15762.

