## Supplement Information for "Type I fatty acid synthase (FAS) trapped in the octanoyl-bound state"

### Table of contents:

#### Supplementary Figures:

Figure S1: Purification and crystallization of the KS-MAT didomain

Figure S2: Validation of ligand placement in the MAT domain

Figure S3: Dynamics of the MAT domain is determined by flexible subdomain linkers

Figure S4: Ramachandran plots for crucial residues in both subdomain linkers

Figure S5: Anisotropic movement of C-alpha atoms in the ferredoxin-like subdomain

Figure S6: Rotational freedom of residue R606

Figure S7: Stability of select KS-MAT variants

Figure S8: Global Michaelis-Menten fit of KS-mediated transacylation data

#### Supplementary Tables

Table S1: Dimerization interface of the KS domain

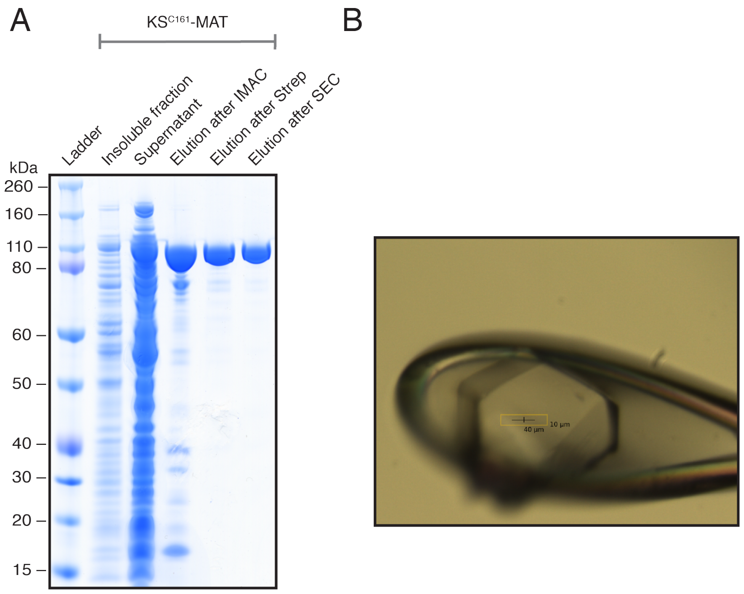


**Figure S1: Purification and crystallization of the KS-MAT didomain.** (A) SDS-PAGE (NuPAGE 4-12 % Bis-Tris) of the purification strategy of the KS-MAT didomain. A tandem purification using Ni-chelating and Strep-Tactin affinity chromatography was followed by size exclusion. (B) Photograph of the octanoyl-CoA soaked crystal within a nylon loop at the synchrotron.

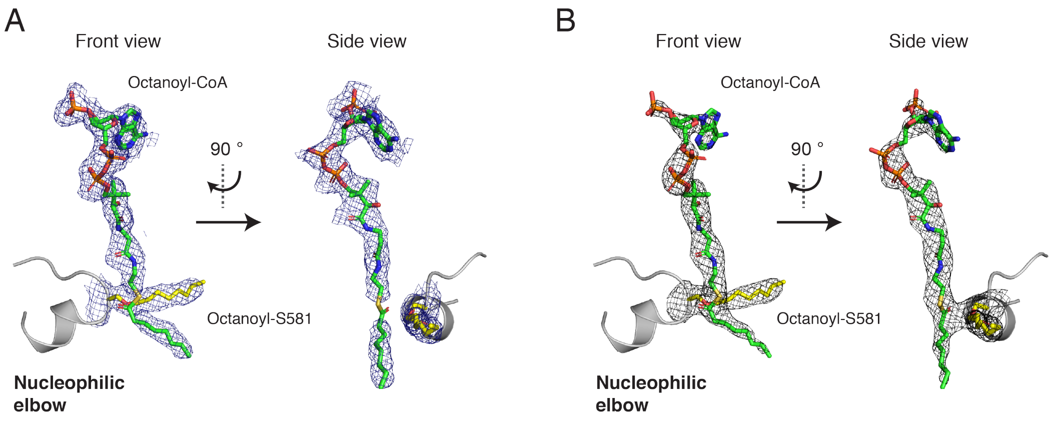


**Figure S2: Validation of ligand placement in the MAT domain.** Octanoyl-CoA and the covalent bound octanoyl-S581 were placed based on unbiased electron density maps. The FEM map (A) is shown in blue and the Polder map (B) in black. Contour levels are at 3 $\sigma$.

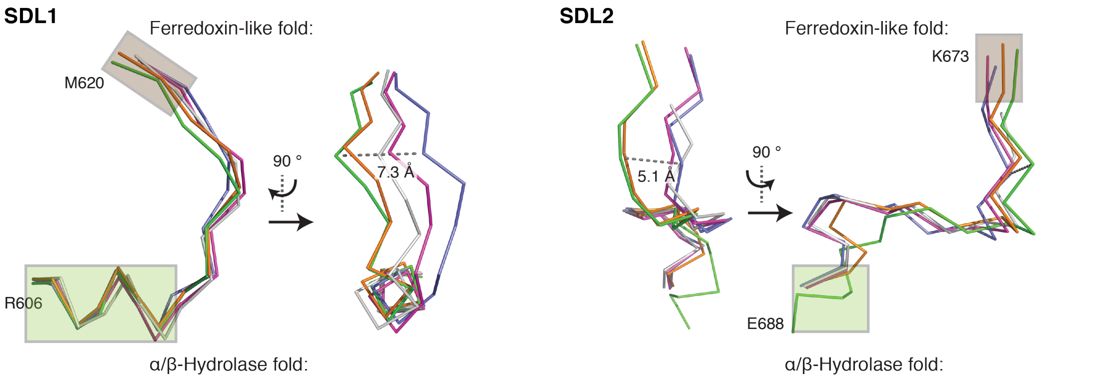


**Figure S3: Dynamics of the MAT domain is determined by flexible subdomain linkers.** SDL1 (612-617, left panel) and SDL2 (675-684, right panel) are shown as ribbons after α/β-hydrolase fold based alignments. Both linker stretches are shown from two different perspectives rotated by 90°. Chain A (blue), chain C (grey) and chain D (green) from the octanoyl-CoA soaked crystal (PDB code 6rop), malonyl-bound (orange) (PDB code 5my0; chain D) and apo human MAT (purple) (PDB code 3hhd; chain A) were used. Green and brown rectangles indicate secondary structure elements of the α/β-hydrolase- and ferredoxin-like subdomains, respectively.

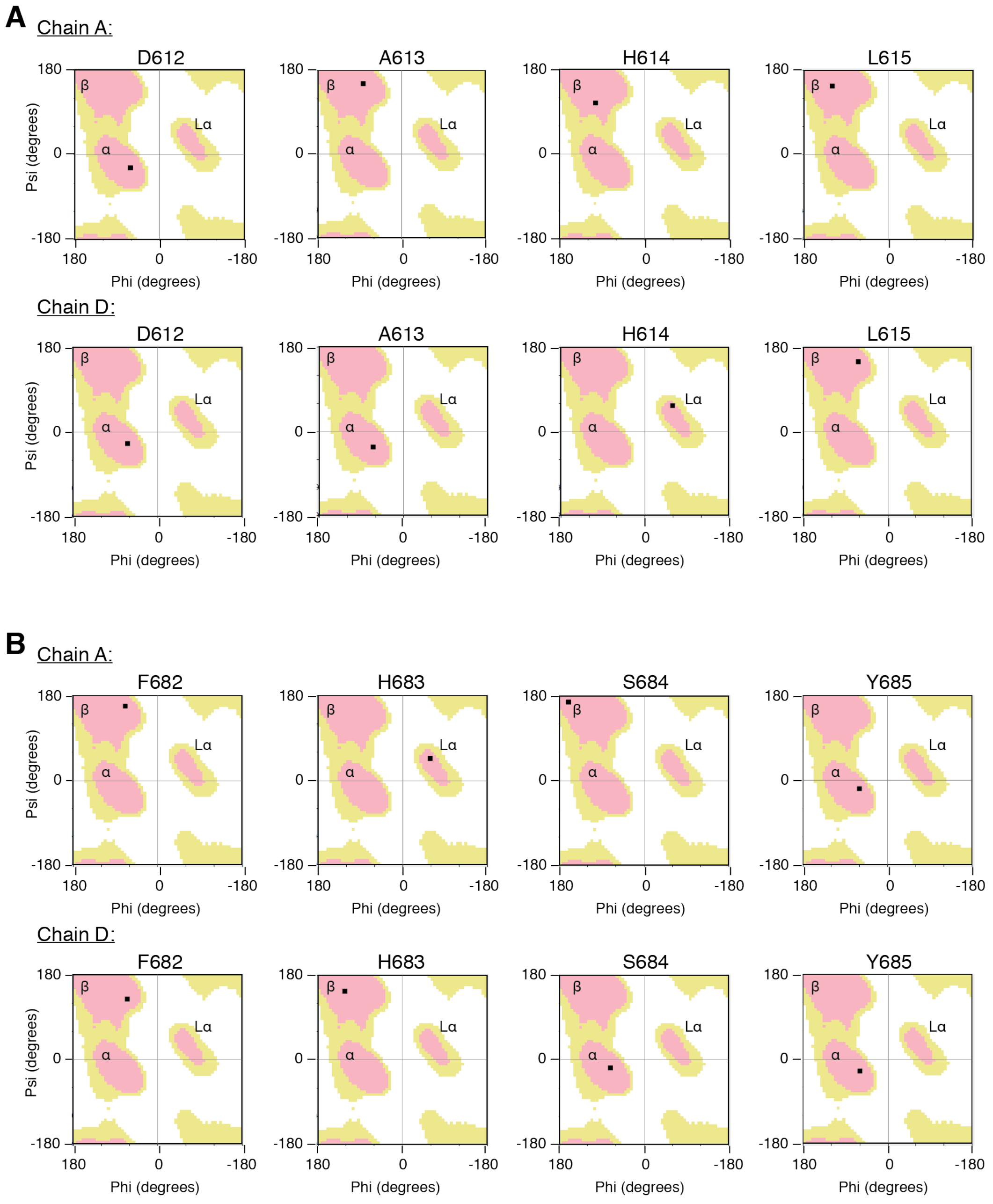


**Figure S4: Ramachandran plots for crucial residues in both subdomain linkers.** Significant changes in dihedral angles were seen for residues in linker 1 (A) 612-615 and linker 2 (B) 682-685 leading to changes to different allowed regions. Plots were created in coot. Used abbreviations indicate $\alpha$ – right-handed helical region, L$\alpha$ – left-handed helical region and $\beta$ – $\beta$-sheet region.

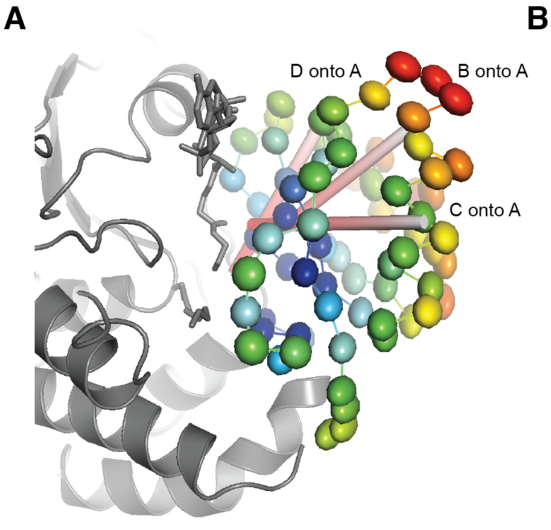


**Figure S5: Anisotropic movement of C-alpha atoms in the ferredoxin-like subdomain.** The anisotropic movement, as derived from the TLS tensors, is depicted by thermal ellipsoids colored by B-values (blue – low values; and red – high values) for the ferredoxin-like fold (616-684) region of chain A (A) and chain D (B). Rotation axes derived from superposition of ferredoxin-like fold (616-684) of chains B (A), C (B), and D (C) onto A (D) are shown in red-white cylindrical bars. For orientation, the rest of the KS-MAT chain is shown in grey.

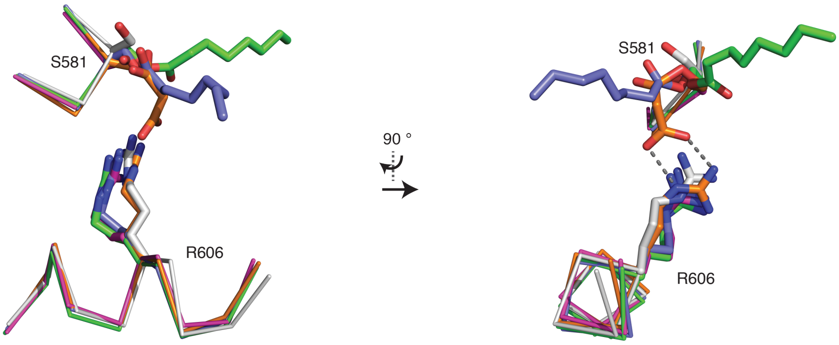


**Figure S6: Rotational freedom of residue R606.** α/β-Hydrolase based superposition (BB of 488-615) of chain A (blue), chain C (white) and chain D (green) from octanoyl-CoA soaked crystals (PDB code 6rop) with the malonyl-bound structure (orange) (PDB code 5my0; chain D) and of porcine FAS (purple) (PDB code 2vz9; chain A). Important residues S581 and R606 are shown in sticks with covalent modifications of the serine represented also in sticks. For clarity residue stretch 580-583 and 601-610 are depicted as ribbons. R606 in octanoyl-bound active sites adopts the same conformation as R606 in the porcine FAS, whereas R606 in chain C (unbound) possesses the rotameric state of the unbound active sites previously found in human KS-MAT (3hhd).[64]

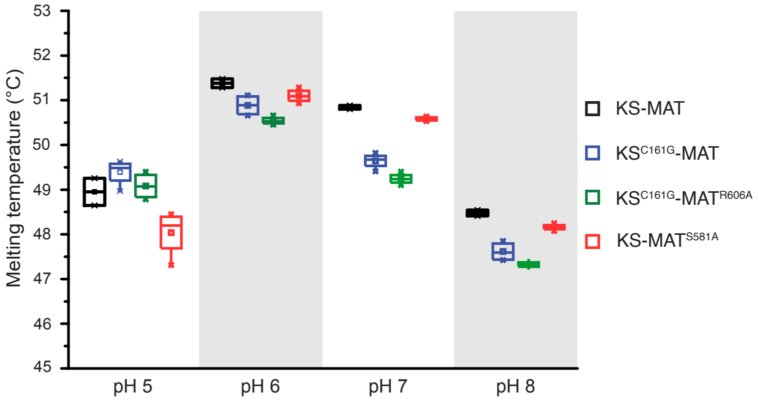


**Figure S7**: **Stability of select KS-MAT variants.** Melting temperatures were determined by a thermal shift assay as described in the Methods section. The four constructs were tested in phosphate buffers at different pH value. Four replicates are shown as dot plot.

A

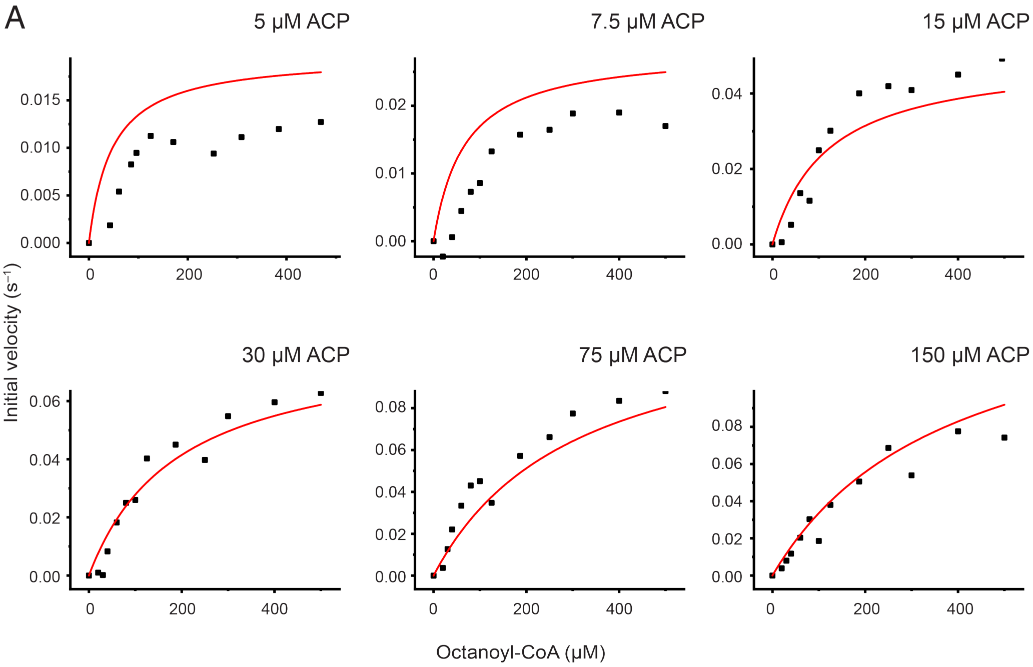


B

| Substrate | $K_m^{X\text{-}CoA}$ (μM) | $K_m^{ACP}$ (μM) | $k_{cat}$ ($s^{-1}$) |
|---|---|---|---|
| C8-CoA | 511 ± 135 | 50 ± 12 | 0.22 ± 0.04 |
| C14-CoA | 231 ± 41 | 7 ± 2.5 | 0.07 ± 0.006 |

**Figure S8: Global Michaelis-Menten fit of KS-mediated transacylation data.** (A) Initial velocities were plotted against octanoyl-CoA (C8-CoA) concentrations at six fixed ACP concentrations. Data were fit globally with the Michaelis-Menten equation assuming a ping-pong bi-bi mechanism. (B) Absolute kinetic parameter derived from the respective global fits for C8- and C14-CoA, respectively. No parameters constraints were set for the fit function.

**Table S1**: Dimerization interface of the KS domain

| Interface | Residues per side | | Solvent-accessible interface area | | Solvent energy gain per side | Hydrogen bonds | Salt bridges |
|---|---|---|---|---|---|---|---|
| | # | % | Å$^2$ | % | kcal/mol | | |
| KS chain A | 73 | 8.6 | 2583 | 8.1 | –14.9 | 38 | 0 |
| KS chain B | 71 | 8.5 | 2578 | 8.0 | –14.8 | | 0 |
| KS chain C | 72 | 8.5 | 2586 | 7.9 | –14.5 | 34 | 0 |
| KS chain D | 75 | 8.8 | 2582 | 8.1 | –15.0 | | 0 |
